# Chimeric deubiquitinase engineering reveals structural basis for specific inhibition of USP30 and a framework for DUB ligandability

**DOI:** 10.1101/2024.09.22.613429

**Authors:** Nafizul Haque Kazi, Nikolas Klink, Kai Gallant, Gian-Marvin Kipka, Malte Gersch

**Affiliations:** Max Planck Institute of Molecular Physiology, Chemical Genomics Center, Dortmund, Germany; TU Dortmund University, Department of Chemistry and Chemical Biology, Dortmund, Germany

## Abstract

The mitochondrial deubiquitinase USP30 negatively regulates Pink1/Parkin-driven mitophagy. Whether enhanced mitochondrial quality control through inhibition of USP30 can protect dopaminergic neurons is currently explored in a clinical trial for Parkinson’s disease. However, the molecular basis for specific inhibition of USP30 by small molecules has remained elusive. Here, we report the crystal structure of human USP30 in complex with a specific inhibitor, enabled by chimeric protein engineering. Our study uncovers how the inhibitor extends into a cryptic pocket facilitated by a compound-induced conformation of the USP30 switching loop. Our work underscores the potential of exploring induced pockets and conformational dynamics to obtain specific deubiquitinase inhibitors and identifies underlying USP30-specific residues. More broadly, we delineate a conceptual framework for specific USP deubiquitinase inhibition based on a common ligandability hotspot in the Leu73-Ubiquitin binding site and on diverse compound extensions. Collectively, our work establishes a generalizable chimeric protein engineering strategy to aid deubiquitinase crystallization and enables structure-based drug design with relevance to neurodegeneration.

## INTRODUCTION

Parkinson’s disease (PD) is a prevalent neurodegenerative disorder, characterized by the progressive loss of dopaminergic neurons in the substantia nigra. A hereditary early-onset form of the disease, termed autosomal-recessive juvenile Parkinsonism, accounts for up to 10% of all cases and has been linked to somatic mutations in genes encoding the Pink1 kinase and the E3 ligase Parkin among others.^1^ Subsequent investigation of these proteins has guided the discovery of a selective autophagy mechanism for mitochondria termed mitophagy and uncovered the central role of dysfunctional mitochondrial quality control in neurons for PD pathogenesis.^2^ In this pathway, Pink1 is stabilized on mitochondria with reduced membrane potential to phosphorylate Ubiquitin conjugated to outer mitochondrial membrane proteins.^3^ This in turn leads to the recruitment and activation of Parkin, which together with the autophagy machinery mediates the lysosomal degradation of damaged portions of the mitochondrial network.^4-6^ Importantly, mitochondrial dysfunction has been strongly implicated in the etiology of both idiopathic and genetic forms of PD.^7^

The post-translational modification of mitochondrial proteins with Ubiquitin thus plays central roles by providing a substrate for Pink1 as well as by mediating autophagy.^8,9^ Consequently, the mitochondrial deubiquitinase USP30 has emerged as a critical negative regulator of mitochondrial quality control due to its ability to remove Ubiquitin from a subset of mitochondrial outer membrane proteins (**Fig. 1a**).^10-15^ Notably, in addition to Pink1/Parkin-driven mitophagy^11,16^, USP30 also negatively regulates basal, i.e. Pink1-dependent but Parkin-independent mitochondrial turnover.^17,18^ Moreover, USP30 regulates mitochondrial protein import^19^, mitochondrial morphology^20^, mitochondrial abundance^21^, apoptotic cell death^22^, and pexophagy^17,23^.

**Figure 1.**
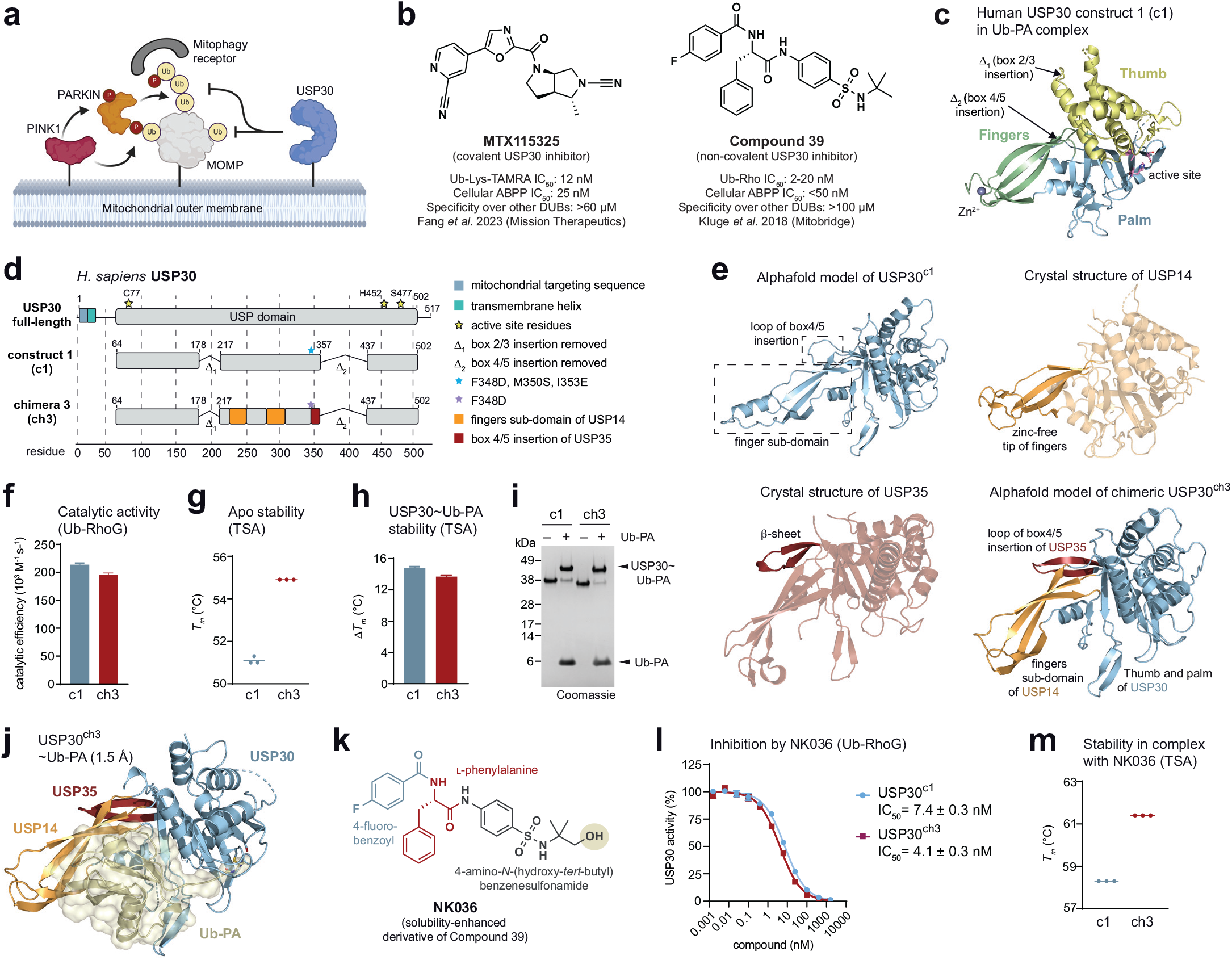
Design, characterization, and inhibition of chimeric USP30. **a**, USP30 antagonizes Pink1/Parkin-mediated mitophagy. **b**, Chemical structures of a covalent and a non-covalent USP30 inhibitor. Inhibition of USP30 to enhance mitochondrial quality control is explored clinically as therapeutic strategy for Parkinson’s and kidney diseases. **c**, Crystal structure of human USP30 obtained with a previously engineered construct as Ub-PA complex (PDB: 5OHK). USP subdomains are shown in different colors. **d**, Architecture of full-length human USP30, previously used construct c1 (named c13 in ^32^) and here described USP30 chimera ch3. See Supplementary Fig. 2 for other chimeras. **e**, AlphaFold-predicted model of USP30^c1^, crystal structures of the catalytic domains of USP14 (PDB: 2AYN) and USP35 (PDB: 5TXK), and AlphaFold-predicted model of USP30 chimera ch3. Regions used for grafting are shown in corresponding colors. **f**, Catalytic efficiencies of indicated USP30 constructs, determined from Ubiquitin-RhoG (Ub-RhoG) cleavage assays. See Supplementary Fig. 3 for raw data. Mean ± s.e.m. **g**, Stability assessment of USP30 constructs in their apo states derived from thermal shift assays. *Tm*, protein melting temperature. **h**, Changes in protein stability upon binding to the Ubiquitin probe Ub-PA. Mean ± s.d. (N=3). **i**, Gel-based Ub-PA binding assay. **j**, Crystal structure of USP30^ch3^∼Ub-PA. See Table 1 for statistics. **k**, Structure of NK036, a solubility-enhanced derivative of Compound 39. **l**, Inhibitory potencies of NK036 for indicated USP30 constructs. IC50 values are given as mean ± s.e.m. **m**, Protein stability of indicated USP30 constructs in the presence of NK036.

A fast-growing body of evidence suggests that USP30 is a highly promising drug target for Parkinson’s disease as its inhibition can protect dopaminergic neurons from α-synuclein-associated toxicity through increased levels of mitophagy.^11,17,24^ This hypothesis has been underscored by studies in neuronal cell lines as well as in flies and in mice.^18,24-28^ Inhibition of USP30 is also being explored as a therapeutic strategy in acute kidney injury due to links to mitochondrial dysfunction.^29^ Several scaffolds of small molecule USP30 inhibitors have been discovered (**Supplementary Fig. 1**),^30,31^ covering covalent and non-covalent modes of inhibition. Two structurally undisclosed compounds have recently been advanced into clinical trials by Mission Therapeutics.

The USP30 inhibitor Compound 39 (**Fig. 1b**) displays IC_50_ values against recombinant USP30 of 2 to 20 nM and cellular target engagement in the 10-50 nM range.^27,33,34^ Moreover, the compound features pronounced specificity for USP30 both in cells and in vitro, with no other DUB being inhibited at 100 µM in a panel of recombinant enzymes – exceeding even the most specific covalent compound.^24,27,34^ Compound 39 with a benzenesulfonamide scaffold^33,35^ as well as the related naphthylsulfonamide MF-094 have been benchmarked in a range of cellular mitophagy assays.^27,34,36,37^ A “pseudo-covalent” binding mode has been proposed due to an exceptionally slow off-rate and compound binding within the catalytic domain has been assessed by hydrogen-deuterium exchange mass spectrometry (HDX-MS).^34^ However, how such outstanding potency and specificity are achieved on the molecular level for any USP30 inhibitor scaffold is currently unknown which has been hindering the acceleration of USP30 inhibitor development. This is likely related to the high flexibility and comparably poor crystallizability of the human USP30 protein for which previously multiple rounds of construct optimization were required for structural characterization in Ubiquitin-bound states (**Fig. 1c)**.^32^

Here, we disclose a generalizable protein engineering strategy based on Ubiquitin-specific protease (USP) deubiquitinase (DUB) chimeras. By grafting structural elements of well crystallizable human USP family members onto the periphery of the human USP30 protein, we generated stabilized and crystallizable protein constructs which retain DUB activity and the propensity to compound inhibition. By solving a high-resolution crystal structure of a suitably optimized USP30 construct in complex with a solubility-enhanced benzenesulfonamide inhibitor, we reveal the molecular basis for the highly potent and specific inhibition of USP30 by small molecules. This method not only elucidated the unique binding mode of this inhibitor class but also suggests a general strategy to enhance the crystallizability of other USP DUBs. We expand our analysis into a conceptual framework for specific USP deubiquitinase inhibition which is based on a common ligandability hotspot in the Leu73-Ubiquitin binding site and diverse compound extensions. Collectively, our work opens new avenues for the structure-based drug design of DUB inhibitors and enables the rational optimization of therapeutics targeting neurodegeneration.

## RESULTS

### Design and characterization of chimeric USP30 constructs

Our work to understand the molecular basis of USP30 inhibition started with an engineered construct of human USP30 (c1), for which crystal structures of Ubiquitin-bound complexes were previously obtained.^32^ This construct contains the three subdomains of human USP30 (palm, thumb and fingers), but lacks two largely unstructured sequence insertions (**Fig. 1c-d, Supplementary Fig. 2a**).^38,39^ However, despite extensive co-crystallization screening with several potent small molecule USP30 inhibitors, no initial crystals were obtained. We hence set out to enhance the crystallizability of the USP30 protein through further construct optimization. We were guided in our approach by a systematic review of all crystal structures of human USP DUBs deposited in the protein data bank (PDB) (**Supplementary Table 1**). Of the 55 human USP enzymes, crystal structures of catalytic domains are available for 21 of these (38%).^40,41^ Notably, 8 of the 55 USP enzymes lack zinc-coordinating cysteine and histidine residues at the tip of their fingers subdomain. Within this subset, 6 of the 8 enzymes have been crystallized (75%) (**Supplementary Fig. 2b**).^39^ This discrepancy is even more pronounced, when focusing on the 14 human USP DUBs for which apo or inhibitor-bound structures were reported (25%). These include the same 6 DUBs which do not bind zinc (**Supplementary Fig. 2b**). These data suggest that USP DUBs lacking a zinc in their fingers subdomain may have a higher propensity for crystallization. A focus on the fingers region for construct optimization was further motivated by (i.) previous HDX-MS analysis suggesting enhanced flexibility compared to other parts of the protein and (ii.) HDX-MS-based mapping of the Compound 39 binding site into the palm and thumb subdomains.^32,34^

We envisaged that the generation of chimeric USP30 catalytic domains, in which the USP30 fingers subdomain is replaced by sequences of equivalent regions in other USP DUBs, may facilitate crystallization (**Fig. 1d-e**). We focused on the fingers domains of USP7 and USP14, which lack zinc ions and have both been crystallized in multiple apo and inhibitor-bound forms (**Supplementary Fig. 2c-f**).^42-48^ We also included the USP DUB CYLD which features truncated, zinc-free fingers (**Supplementary Fig. 2g**),^49,50^ and also designed construct 2 (c2) in which the entire fingers were replaced by Gly-Ser linkers (**Supplementary Fig. 2h**). In addition, we noticed that the loop of the box 4/5 insertion deletion features high flexibility in HDX-MS and elevated *B*-factors in Ubiquitin-bound complexes. We planned a replacement by the equivalent region of USP35, as USP35 is the only DUB featuring secondary structure in this region (a short, antiparallel β-sheet, **Supplementary Fig. 2f**).^51^

In an iterative process, we explored 15 chimeric constructs starting with boundary design by structure superposition, design validation by Alphafold-based modeling (**Supplementary Fig. 2h-l**), cloning (**Supplementary Fig. 2m**), protein purification, and biophysical characterization (**Supplementary Fig. 3a-e**). This process is illustrated with four diverse chimeras: Chimera 1 (USP30^ch1^) features the fingers of USP7, chimera 2 (USP30^ch2^) features the fingers of USP14, chimera 3 (USP30^ch3^) features the fingers of USP14 and the box 4/5 insertion of USP35, and chimera 4 (USP30^ch4^) features the fingers of CYLD (**Supplementary Fig. 2i-l)**. All proteins were expressed, purified, and their stability was assessed in thermal shift assays (TSA). While inclusion of the USP14 and CYLD fingers did not alter protein stability, USP7 fingers destabilized the chimeric protein, whereas inclusion of the structured box 4/5 insertion of USP35 increased protein stability (**Supplementary Fig. 3a**). Chimeras 1-3 showed complete binding to the Ubiquitin-probe Ub-PA (**Supplementary Fig. 3b**), which correlated with protein stabilization (**Supplementary Fig. 3c**), whereas this was not the case for constructs with truncated fingers. Consistently, chimeras 1-3 showed high catalytic activity towards the fluorogenic substrate Ubiquitin-RhoG (Ub-RhoG, **Supplementary Fig. 3d-e**), whereas USP30^ch4^ and construct 2 were virtually inactive. These results are consistent with the large protein interaction surface contributed by the fingers for Ubiquitin recognition. Importantly, chimeras 1-3 retained their propensity to be inhibited by Compound 39 with IC_50_ values between 0.3 and 0.7 nM compared to 0.8 nM of construct 1 (**Supplementary Fig. 3f-g**). To assess inhibitor binding in all proteins, including the catalytically inactive chimeras, we measured inhibitor-induced changes in protein stability. The presence of Compound 39 increased protein stabilities between 7 and 9°C for all samples, which demonstrates that all USP30 constructs retain affinity for the benzenesulfonamide scaffold (**Supplementary Fig. 3h**).

### Design validation and inhibition of a chimeric USP30-(USP14-USP35) construct

Upon surveying the collected data (**Supplementary Table 2**), we decided to focus on USP30^ch3^. This construct containing the USP14 fingers and the USP35 box 4/5 insertion (**Fig. 1d-e**) features unaltered catalytic activity (**Fig. 1f**), displays an increased protein stability by approx. 4°C (**Fig. 1g**) and reacts readily with Ub-PA (**Fig. 1h-i**). To validate our design, we solved the crystal structure of the covalent USP30^ch3^∼Ub-PA complex (**Fig. 1j, Table 1**). Both a rather high rate of initial crystal hits in coarse-screening plates and the high resolution of 1.5 Å without crystallization fine-screening supported the hypothesis of increased crystallizability of this chimeric protein. Inspection of the electron density revealed that the boundary design allowed seamless chimeric sequence transitions and did not perturb the USP fold (**Supplementary Fig. 4a**). Consistently, the structure showed a highly similar arrangement compared to Ub-PA complexes of USP30, USP14 and USP35 (**Supplementary Fig. 4b**).

**Table 1.**
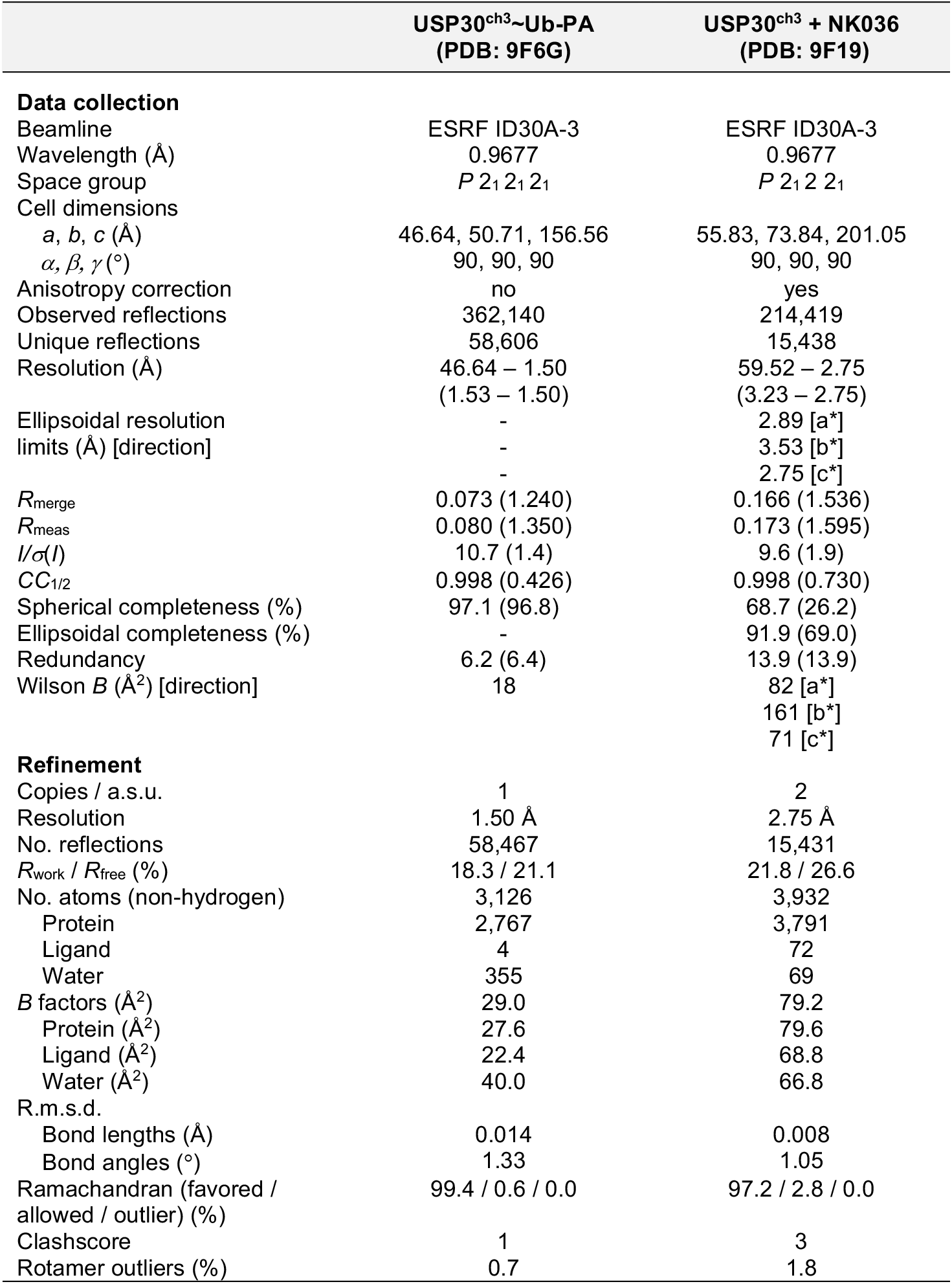
Data collection and refinement statistics.

During the initial crystallization experiments, we noticed that Compound 39 possesses poor solubility in aqueous buffers. We therefore synthesized NK036 as solubility-enhanced derivative (**Fig. 1k**). This compound shares the 4-fluorobenzoyl group, the central L-phenylalanine and the benzenesulfonamide with Compound 39 and features an additional hydroxyl group at the *tert*-butyl group. While this addition decreased potency by about one order of magnitude (with IC_50_ values between 4 and 7 nM, **Supplementary Fig. 3f-g**), it improved NK036 solubility in aqueous buffers. Importantly, NK036 inhibits USP30^c1^ and USP30^ch3^ to a similar degree (**Fig. 1l**) and increases protein stability to above 61°C (**Fig. 1m**), with the same relative stabilization of all constructs (**Supplementary Fig. 3h**). These data underscore the suitability of the stabilized USP30^ch3^ construct and of NK036 as potent USP30 inhibitor for co-crystallization studies.

### Crystal structure of USP30^ch3^ in complex with a non-covalent inhibitor

The structure of USP30^ch3^ in complex with NK036 was solved to 2.75 Å following crystal optimization through fine-screening and additive screening as well as data optimization through multi-crystal averaging and anisotropic scaling (**Fig. 2a, Table 1, Supplementary Table 3**, see the methods section for details). The asymmetric unit contained two copies of the protein-ligand complex. These could be superimposed with a Cα-RMSD of 0.4 Å (**Supplementary Fig. 5a**) and showed near identical ligand geometries (**Supplementary Fig. 5b**). Both chains displayed clear electron density which allowed unambiguous positioning of the inhibitor (**Fig. 2b** and **Supplementary Fig. 5c**). Chain A contained more ordered residues near the fingers subdomain, which were partially disordered in chain B, whereas conversely chain B showed ordered loops on the opposite side near the thumb subdomain.

**Figure 2.**
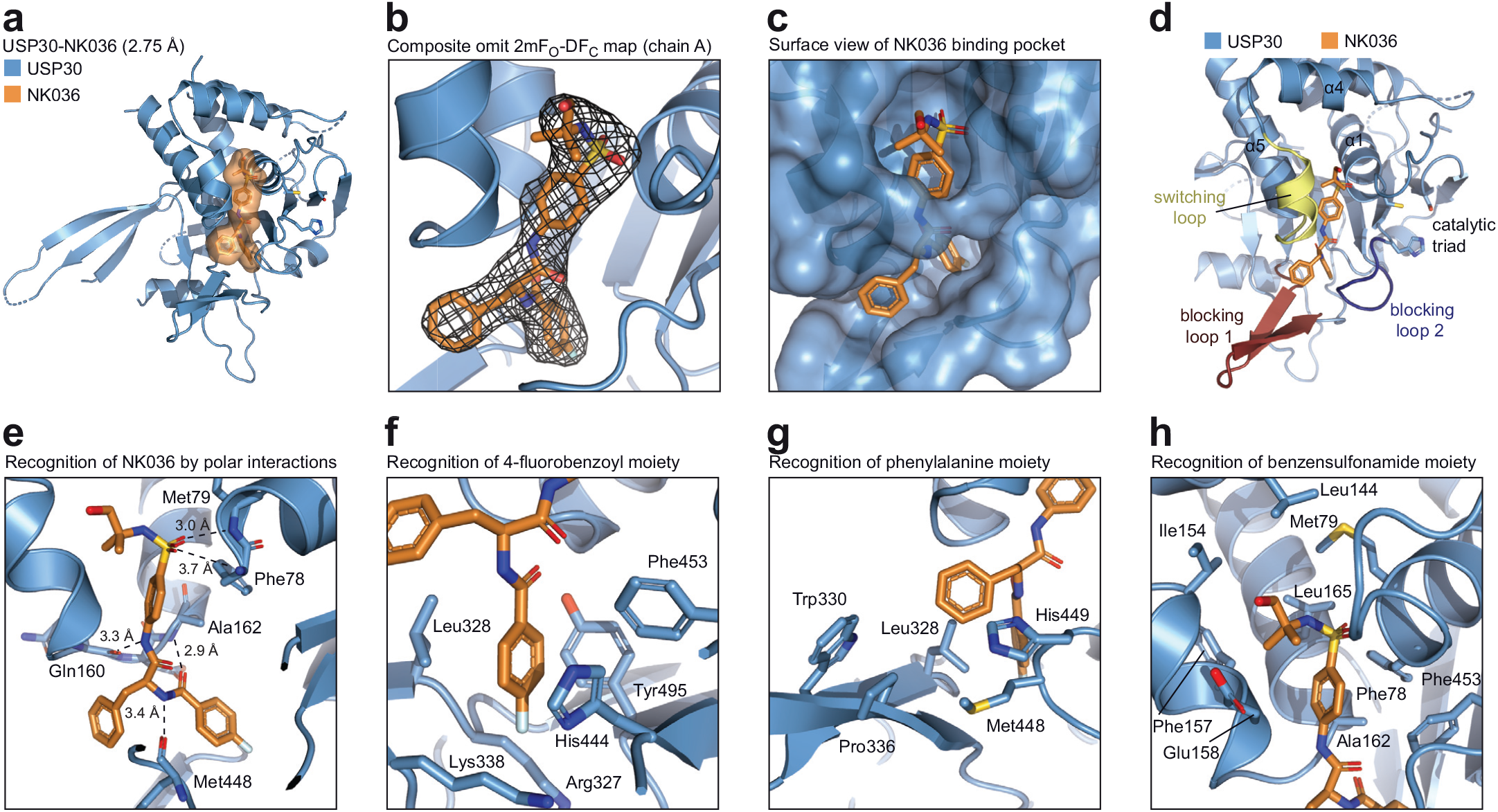
Structure of USP30 in complex with NK036. **a**, Cartoon representation of the crystal structure of USP30^ch3^ bound to NK036. The compound is shown under an orange surface. **b**, Composite omit electron density map of NK036 in chain A (2mFO-DFC, contoured at 1σ, covering all atoms of the compound). **c**, Structure as in A with surface representation of USP30. **d**, Compound binding site highlighting typical USP regions involved in binding to NK036. **e**, Close-up view of the compound binding site highlighting key residues involved in H-bonding. **f**, Close-up view of the USP30 hydrophobic patch engaging the fluorobenzoyl moiety of the compound. **g**, Close-up view of hydrophobic interactions of the phenylalanine group of the compound. **h**, Close-up view of the benzenesulfonamide moiety of the compound engaged by USP30 residues.

These differences could be attributed to crystal contacts and these regions are far away from the ligand binding site. Thus, chain A was used for all further analysis. The ligand was surrounded only by USP30-encoded side chains as well as secondary structures generated by USP30 residues, with chimeric portions of the sequence spatially separated (**Supplementary Fig. 5d**). This indicates that the chimeric engineering in USP30^ch3^ does not prevent the deduction of a *bona fide* USP30 inhibition mechanism.

NK036 was found to be engaged by both the palm and thumb subdomains (**Supplementary Fig. 5e**) and extensively surrounded by protein residues (**Fig. 2c** and **Supplementary Fig. 5f**). The phenylsulfonyl and *tert*-butyl moieties are engaged by the thumb subdomain, being buttressed between the switching loop, the α5 helix and the α1 helix. The phenylalanine and fluorophenyl moieties of the ligand are mainly surrounded by blocking loops 1 and 2 within the palm region (**Fig. 2d**). The two central amide bonds of the ligand are contacted through a total of three hydrogen bonds by the protein (**Fig. 2e**): The backbone carbonyl group of Gln160 engages the anilinic NH proton, the carbonyl group of Met448 contacts the other amidic NH proton, and the carbonyl group of the fluorobenzoyl moiety engages the Ala162 nitrogen. These polar interactions give rise to a star-like tripartite geometry of NK036 with the hydrophobic portions of the ligand extending into three separate areas:

a. The fluorobenzoyl ring binds into a hydrophobic pocket formed by Leu328, Tyr495, Phe453, as well as aliphatic portions of the Arg327 and Lys338 side chains (**Fig. 2f**). While the imidazole ring of His444 binds the fluorophenyl ring through parallel π-π stacking, the Leu328 side chain on the other side of the ring creates a pin to close the pocket towards blocking loop 1.
b. The central phenyl ring of the ligand is engaged by hydrophobic interactions with Leu328, Met448, and parallel π -π stacking with His449 (**Fig. 2g**). Its tip is near Trp330 and Pro336 on blocking loop 1, while not fully occupying the binding groove. This is in line with compounds featuring a larger cyclohexylmethyl group instead of the phenyl ring also displaying potent USP30 inhibition.^33^
c. The benzenesulfonamide is surrounded by Ala162, Phe78, Phe453 and the aliphatic part of the Glu158 side chain (**Fig. 2h**). The sulfonyl is contacted by the Phe78 and Met79 amides (**Fig. 2e**). The *tert*-butyl group is bound in a hydrophobic cleft which is formed towards the top by Met79, Leu165 and Leu144 side chains, and on the side by Ile154 and Phe157. The electron density did not allow positioning of the hydroxyl group of NK036 (also reflected in the difference between chain A and chain B geometries, **Supplementary Fig. 5b**), suggesting that it is not specifically engaged by the protein. This observation in combination with the hydrophobic environment explains the potency (**Fig. 1k-l**) as well as the tolerance of other hydrophobic structural elements at this position within the compound series (**Supplementary Fig. 1**).^33,35^

Overall, the structure reveals extensive contacts of all regions of the inhibitor and rationalizes observed structure-activity relationships. The large interaction surface of 570 Å^2^ and the deep embedding of the ligand into the protein fold explain the very slow off-rate and previously observed pseudo-covalent binding characteristics.^34^ Notably, the regions of the ligand binding site are in perfect agreement with the previous HDX-MS analysis. However, the experimentally determined compound binding mode is distinct from docking models obtained from fitting the compound into Ub-bound geometry of USP30.^34^ This is due to unexpected conformation differences further analyzed below.

### Conformational plasticity of the switching loop and engagement of the Leu73 pocket underly USP30 ligand binding

To understand how binding to NK036 inhibits USP30, we compared our structure to the USP30∼Ub-PA complex (**Fig. 3a-d**). While the catalytic residues of USP30 are not contacted by the inhibitor, we observed that the catalytic triad was not aligned in the inhibitor bound state as the catalytic histidine was flipped out (**Fig. 3d**). The superposition further showed the phenylalanine and fluorobenzoyl moieties of NK036 to occupy the cleft which guides the Ubiquitin C-terminus to the USP30 active site. This is accompanied by small changes in blocking loop 2. This substrate competitive binding mode is facilitated by the fluorobenzoyl group binding into a pocket which is used to recognize the Ubiquitin Leu73 sidechain, with an inhibitor amide taking the place of the Ubiquitin Leu73-Arg74 amide (**Fig. 3e-f**). NK036 thus inhibits USP30 by preventing Ubiquitin binding.

**Figure 3.**
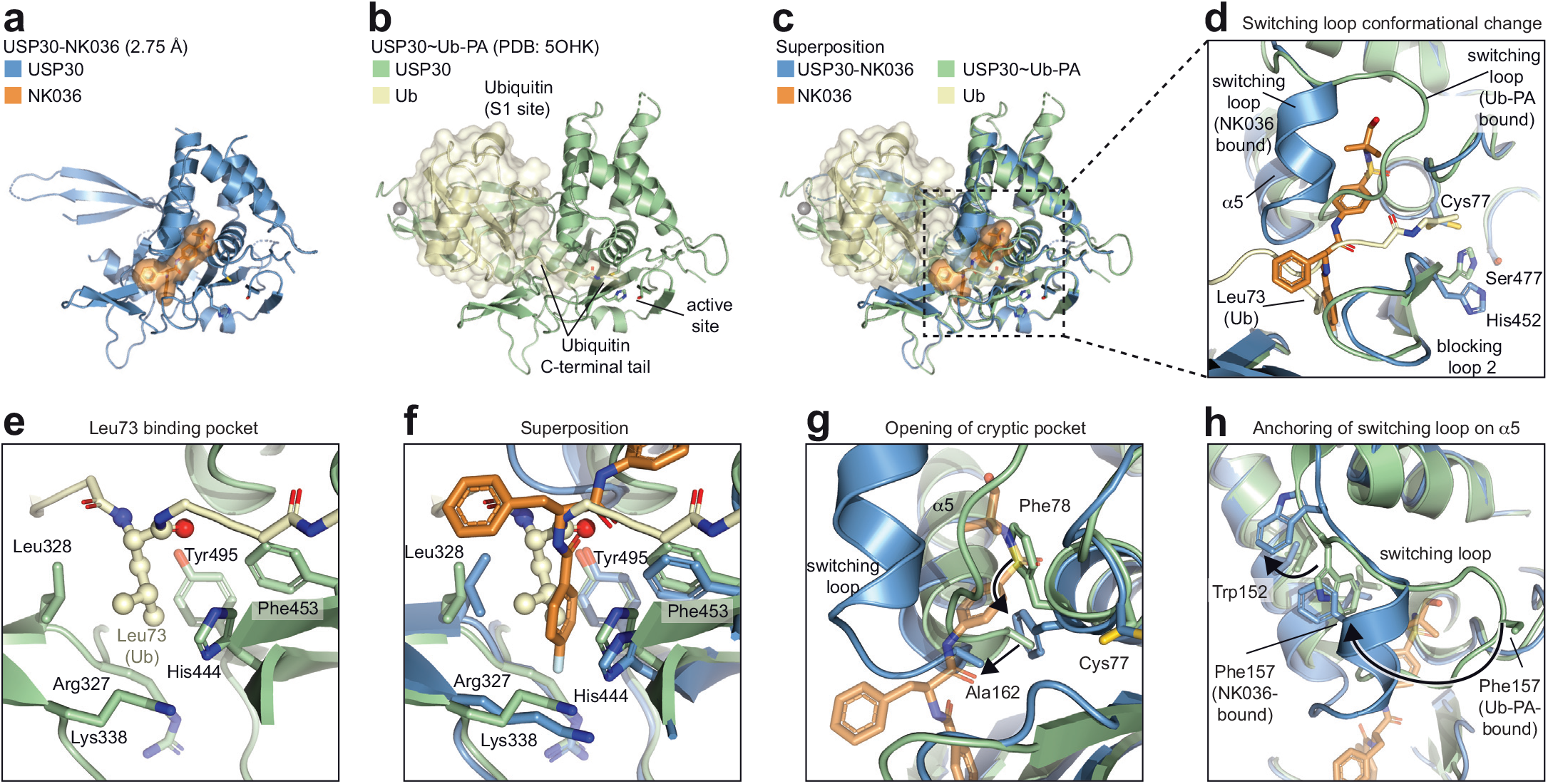
Engagement of the Leu73 pocket and conformational plasticity of the switching loop underlie ligand engagement by USP30. **a**, Cartoon representation of the crystal structure of USP30 bound to NK036. The compound is shown under an orange surface. **b**, Structure of USP30∼Ub-PA complex (PDB: 5OHK). Ubiquitin is shown under a yellow surface. **c**, Superposition of panels a and b. **d**, Close-up view of the compound binding site. Catalytic triad residues of USP30 and Leu73 of Ubiquitin are labeled. The conformational change of the USP30 switching loop is indicated. **e**, Close-up view on the engagement of the Ubiquitin Leu73 side chain by USP30, with residues forming the hydrophobic pocket highlighted. **f**, Superposition of the structures, focused on the Leu73 binding pocket showing its occupation by the fluorobenzoyl group of NK036. **g-h**, Close-up views of the conformational changes of the switching loop, focusing on entry of the cryptic pocket within the thumb subdomain (g) and the anchoring of the switching loop on the α5 helix (h). Putative movements of residues are indicated with arrows. See also **Supplementary Movie 1**, in which the equivalent transition is shown based on the Lys6-diubiquitin-bound structure of USP30 (PDB: 5OHP).^32^

The superposition also revealed large and surprising conformational changes of the switching loop which allow the *N*-*tert*-butyl-benzenesulfonamide group to bind deeply within the thumb subdomain (**Fig. 3d**). This cryptic pocket is not present in previously analyzed Ubiquitin-bound states. We next wanted to understand how this novel conformation is facilitated. Closer inspection of the associated residues revealed the benzenesulfonamide to take the place of Phe78 which moves inwards and takes the position of Ala162. This change in turn pushes the tip of the α5 helix outwards by approx. 3 Å to generate an entry into the pocket (**Fig. 3g**). The switching loop is anchored on the α5 helix by Phe157 which creates a side of the pocket. Notably, Phe157 moves more than 18 Å and takes the place of Trp152, which is pushed out towards the fingers (**Fig. 3h, Supplementary Movie 1**). This surprising conformational change of the switching loop is accompanied by a loop-to-helix transition with residues 154-159 forming a two-turn alpha helix – a conformation not observed for USP30 or any other USP switching loop (**Supplementary Fig. 6a-h**). As there is no apo structure of USP30 available, we cannot rigorously distinguish between inhibitor-stabilized and *de novo* inhibitor-induced states. However, the observations that the switching loop shows very high flexibility by HDX-MS in the apo state and that the observed conformation is incompatible with Ubiquitin binding strongly suggest that this is an inhibitor-induced conformation, which is facilitated by structural plasticity of the switching loop.

This conformational change also represents a second mechanism by which Ubiquitin engagement is prevented in the inhibitor-bound state.

### Molecular basis for specific inhibition of USP30

We next asked why the inhibitor series displays pronounced specificity for USP30 both in vitro and in cells.^27,34^ Analysis of the binding site revealed that many residues contacted by the inhibitor are strictly conserved in many other USP family members.^39^ These include Phe78 and Met79 directly adjacent the catalytic cysteine, His444 on blocking loop 2, Ala162 at the start of α5, and Tyr495. However, two residues stood out which are unique for USP30 within the entire human USP family: Leu328 where other USP DUBs typically feature a phenylalanine, and Phe453 instead of which a tyrosine is common (**Fig. 4a**). Both residues were noted previously during the analysis of atypical Ubiquitin binding by USP30^32^ and are located within the center of the ligand binding site (**Fig. 4b**). Superpositions with structures of other DUBs featuring the canonical USP residues at these positions indicate how these would interfere with ligand binding (**Fig. 4c**). The additional phenolic hydroxyl group of a tyrosine at position 453 would clash with the carbonyl oxygen of a ligand amide, whereas a larger phenylalanine at position 328 would interfere with two of the three hydrophobic pockets due to Leu328’s role as pin between these. This analysis suggested that Leu328 and Phe453 are key specificity factors for USP30 inhibition by benzenesulfonamide inhibitors.

**Figure 4.**
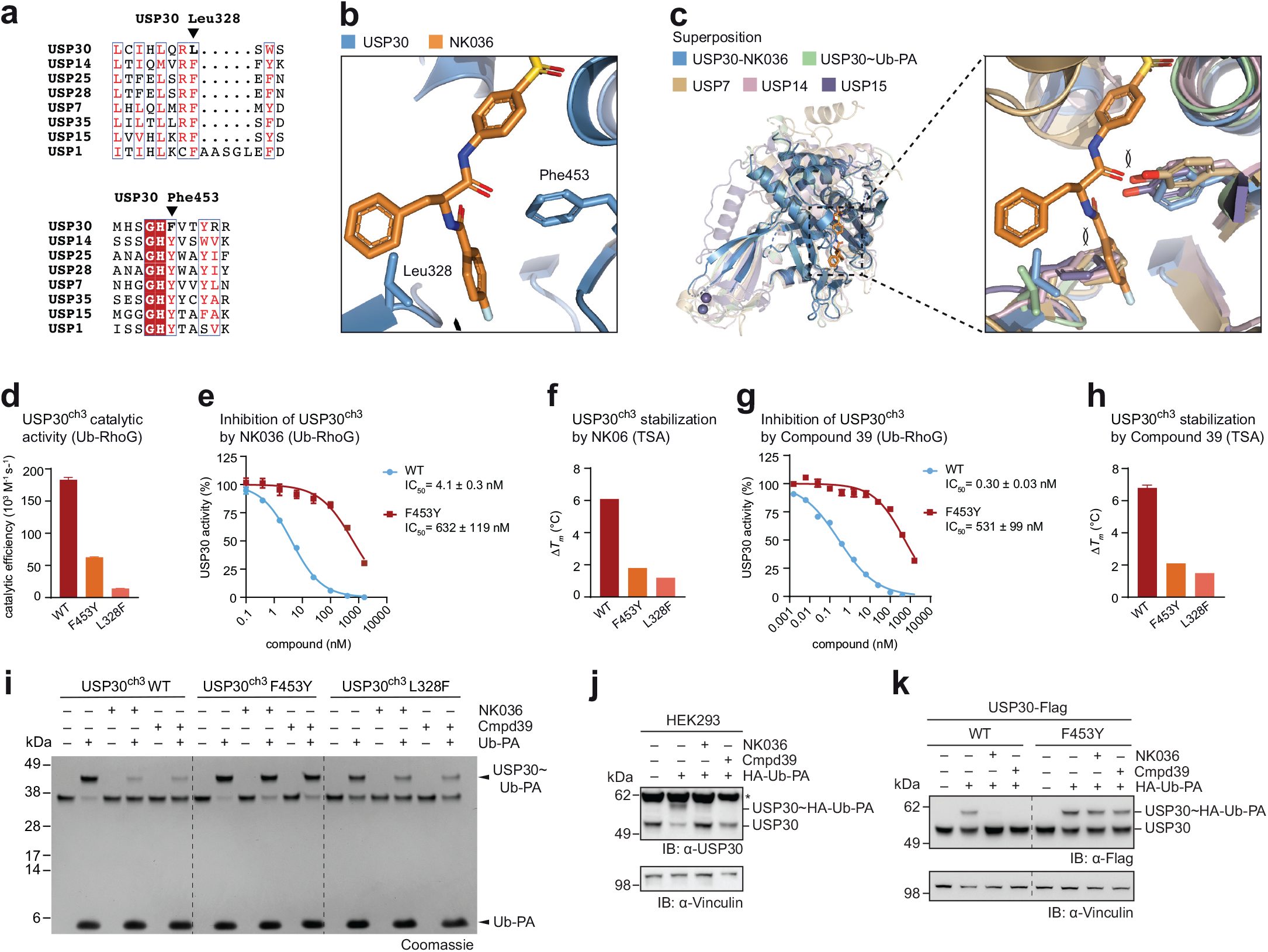
Molecular basis of inhibitor specificity for USP30. **a**, Sequence alignment of indicated human USP DUBs. Arrows indicate the unique Leu328 and Phe453 residues in USP30. **b**, Close-up view of the compound binding site. **c**, Superposition with indicated USP DUB structures in complex with inhibitors (PDB: 5N9R, 6IIN, 6GH9), highlighting how equivalent Phe and Tyr residues in other human USP DUBs interfere with compound binding. **d**, Catalytic activities of indicated wild-type (WT) and mutant USP30 proteins, assessed by Ub-RhoG cleavage. Mean ± s.e.m. **e**, Inhibitory potencies of NK036, pre-incubated with indicated USP30 proteins for 1.5 h, determined from Ub-RhoG cleavage assays. IC50 values are given as mean ± s.e.m. **f**, Protein stability of indicated USP30 proteins in the presence of 20 µM NK036. Δ*Tm* was calculated as *Tm* of the compound-bound sample subtracted from *Tm* of the respective apo protein. Mean ± s.d. (N=3). **g**, Inhibitory potencies of Compound 39, determined as in e. **h**, Protein stability assessment in the presence of Compound 39, determined as in f. **i**, Probe competition assay. USP30 and inhibitors were preincubated, followed by addition of Ub-PA, and analysis of samples by SDS-PAGE and Coomassie staining. **j**, Cellular probe competition assay. HEK293 cells were treated with indicated compounds. Lysates were then incubated with ubiquitin probe where indicated and analyzed by western blot. The asterisk denotes an unspecific band. **k**, Cellular assessment of USP30 inhibition mechanism. C-terminally Flag-tagged USP30 (WT or compound-resistant mutation F453Y) was overexpressed in HEK293 cells. Cells were analyzed as described in panel j.

To experimentally test this hypothesis, we generated F453Y and L328F point-mutated proteins. While the reduced catalytic activity of L328F did not allow enzyme inhibition assays at 0.1 nM enzyme concentration, the F453Y mutant showed reduced but robust cleavage of Ub-RhoG (**Fig. 4d** and **Supplementary Fig. 6i-j**). Substitution of Phe453 by the canonical tyrosine reduced inhibition potency of NK036 by more than two orders of magnitude (**Fig. 4e**), which validates the observed binding mode. Consistently, both mutant proteins were less stabilized by NK036 in thermal shift assays. TSA assays were carried out at a compound concentration (20 µM) above the inhibitory IC_50_, explaining partial protein stabilization (**Fig. 4f**). We also repeated the analysis with Compound 39 with identical results, identifying compound-resistant activity of the F453Y mutant with a three orders of magnitude specificity window (**Fig. 4g-h**). These data demonstrate that two residues which are unique for USP30 within the human USP DUB family and which are in the center of the inhibitor binding site are key to facilitating specific inhibition of USP30.

We next set out to assess this mechanism of USP30 inhibition in a cellular context. To this end, we first optimized a competitive probe labeling assay. While preincubation of inhibitors with wild-type recombinant USP30 largely abrogated labeling with Ub-PA, USP30 carrying the compound-resistant F453Y mutation showed complete labeling in all conditions (**Fig. 4i**). The same trend could be observed for the L328F mutation, however, its reduced probe labeling precluded further analysis. The activity of both Compound 39 and NK036 on endogenous USP30 was then confirmed through a cellular probe-labeling experiment (**Fig. 4j**), consistent with previous results.^27^ Next, we overexpressed Flag-tagged USP30 in HEK293 cells and carried out probe labeling experiments (**Fig. 4k**). Treatment of cells for 2 h with either Compound 39 or NK036 abolished probe labeling of the wild-type protein. In contrast, USP30 carrying the F453Y mutation was equally labeled by the ubiquitin probe when cells were previously treated with inhibitors. These data confirm the compound-resistance of this mutation in the context of the full-length protein and, more broadly, provide cellular validation for the observed mechanism of specific USP30 inhibition by benzenesulfonamides.

### NK036 engages DUB ligandability hotspot in unique manner

To understand how the identified mechanism of specific USP30 inhibition differs from those of other non-covalent USP DUB inhibitors, we analyzed published human USP:ligand structures. These included USP7 in complex with hydroxypiperidine inhibitors (**Fig. 5a**)^46-48^, USP7 in complex with an allosteric inhibitor (**Fig. 5b**)^45^, USP14 bound to IU1-206 (**Fig. 5c**)^44^, USP28 in complex with FT206 (**Fig. 5d**)^53,55^, USP1 in complex with ML323 (**Fig. 5e**)^52^, and USP7 in complex with Compound 23 (**Fig. 5f**)^54^. In addition, we also compared the binding mode to a covalent USP7 inhibitor^46^ as well as to an inhibitor of the USP-fold SARS-CoV PLpro enzyme (**Supplementary Fig. 7a-d**)^56,57^. Superposition of all structures with the obtained USP30-NK036 geometry revealed the benzenesulfonamide moiety of NK036 to engage a previously unexplored pocket within the USP domain (**Fig. 5g-h**).

**Figure 5.**
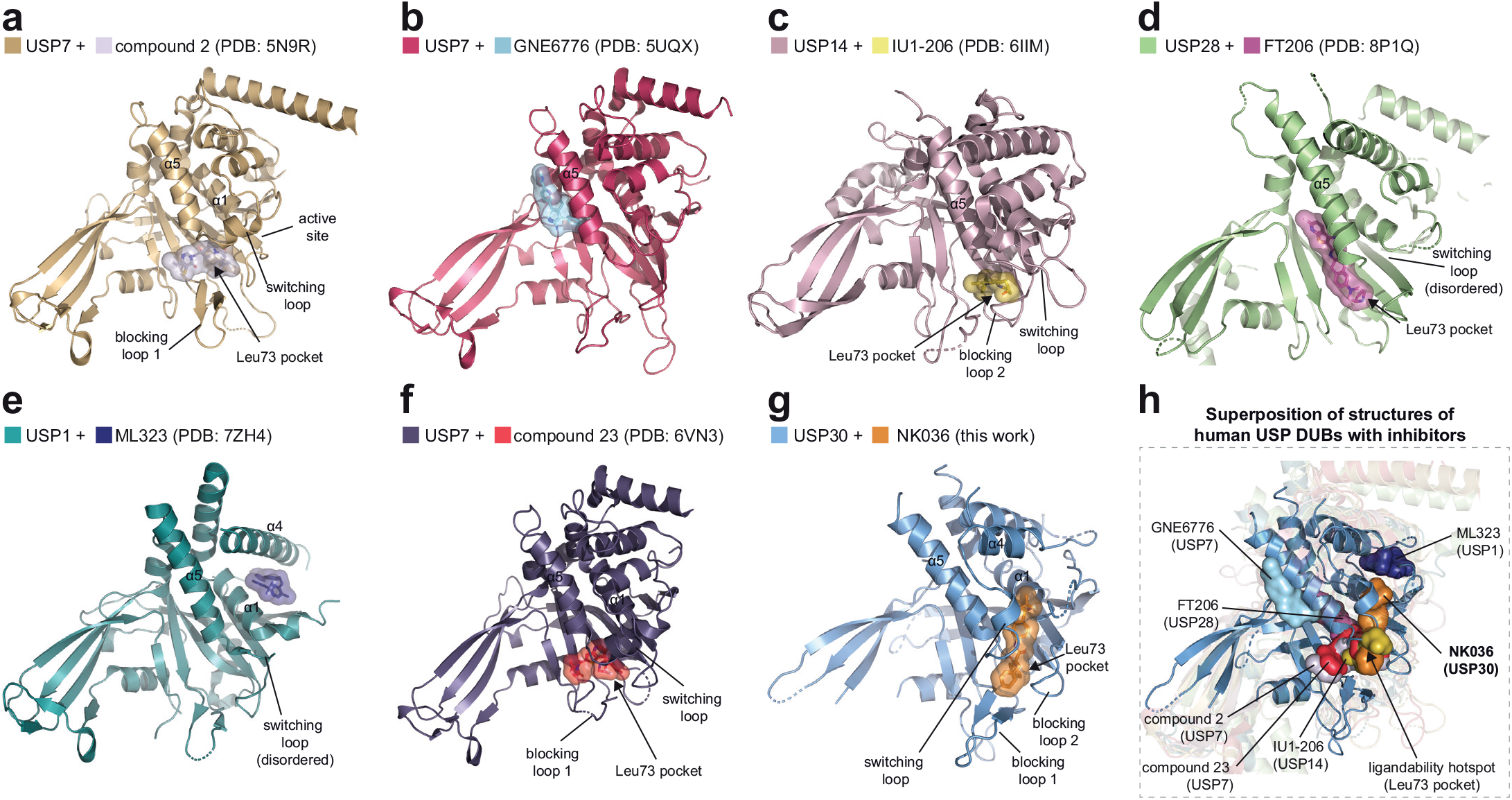
Inhibition of USP30 by NK036 uses a ligandability hotspot and a cryptic pocket, distinct from other USP DUB inhibitors. **a-g**, Cartoon representations of human USP-family deubiquitinase catalytic domains in complex with indicated small molecule inhibitors. Compounds are shown as both sticks and transparent surfaces. Structural elements of DUBs are labeled and PDB codes of structures are given.^44,45,48,52-54^ The Leu73 Ubiquitin binding site is shown with an arrow when engaged by compounds. **h**, Comparison of USP30 inhibition by NK036 to other DUB inhibitors. Superposition of the structure of USP30+NK036 on other structures shown in panels a-f. Compounds are shown as surfaces and are labeled. All USP cartoons except USP30 are semitransparent.

The fluorophenyl and phenylalanine moieties of NK036 overlap with binding modes determined for non-covalent USP7, USP14, and USP28 inhibitors as well as for covalent USP7 and PLpro inhibitors. Closer analysis of the structural superpositions revealed the para-fluorophenyl group of NK036 to engage the Leu73 Ubiquitin binding site of USP30 in the same way as the chemically related para-chlorophenyl- and 3-fluoropyrazole groups of USP7 and USP14 inhibitors, which are otherwise structurally completely unrelated (**Supplementary Fig. 8**).^44,46,54^ We previously identified the molecular basis for specific inhibition of the UCH-family deubiquitinase UCHL1 by covalent cyanamides^58,59^ and in this context proposed the binding pocket of DUBs for the Ubiquitin Leu73 side chain as general DUB ligandability hotspot.^58^ Our analysis reveals that NK036 engages USP30 through precisely this hotspot in a unique manner as it also expands through its benzenesulfonamide moiety into a cryptic pocket in the thumb subdomain, which is generated through conformational plasticity of the switching loop.

## DISCUSSION

Due to the multi-facetted cellular roles of the Ubiquitin system, many novel therapeutic approaches are currently explored by modulating Ubiquitin signaling with small molecules.^60^ Deubiquitinase inhibitors have the potential to amplify Ubiquitin-dependent processes and inhibitors of two DUBs are currently being evaluated in clinical trials. These include compounds targeting the mitochondrial DUB USP30 for kidney disease and Parkinson’s disease to elevate mitochondrial quality control, yet how the specific inhibition of USP30 can be facilitated on the molecular level had remained elusive.^61^ Through chimeric engineering, we here report a structure of human USP30 in complex with a potent inhibitor. The devised constructs will also aid the structural investigation of other USP30 inhibitors, and the obtained structure will enable rational design and discovery of improved compounds while retaining the excellent potency and specificity of the scaffold.

Our investigation reveals the molecular basis for potent and specific inhibition of USP30. NK036 uses its fluorobenzoyl moiety to occupy the unique Leu73 recognition pocket of USP30 and additionally occupies a cryptic pocket within the thumb subdomain with its benzenesulfonamide group (**Fig. 6a-b**). This binding mode is facilitated by a loop-to-helix transition of the switching loop, for which solvent-exposed residues are repurposed to lock the loop in a structured, domain-bound state. This loop shows high sequence variability between human USP DUBs and has previously been discussed in the context of specific USP7 inhibition, where compounds of the hydroxypiperidine scaffold stabilize the inactive apo state of USP7.^46,47^

**Figure 6.**
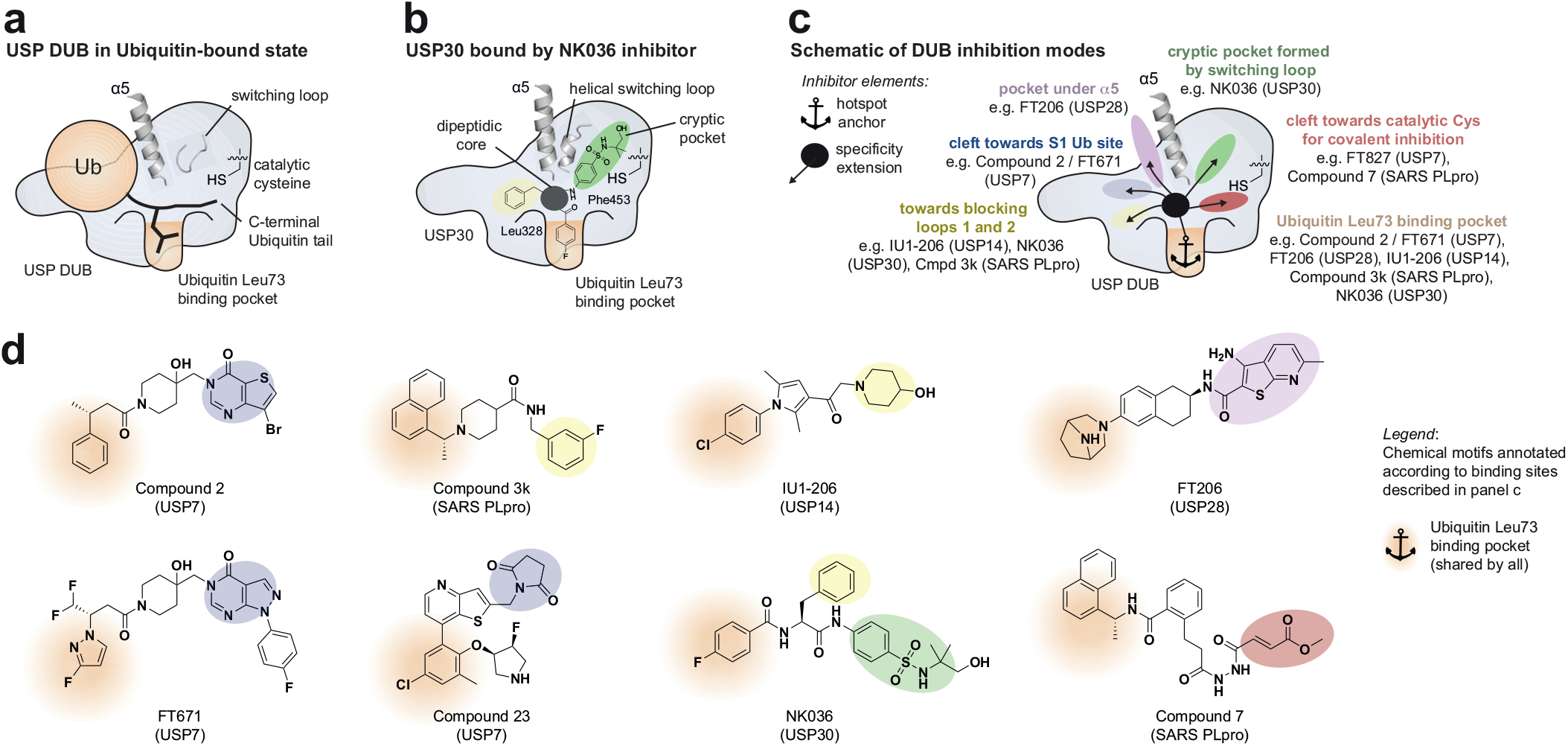
A framework for specific USP deubiquitinase inhibition. **a**, Schematic of a Ubiquitin-bound USP DUB. Structural elements are labeled. **b**, Schematic of USP30 in complex with an inhibitor of the benzenesulfonamide scaffold, occupying the Ubiquitin Leu73 pocket (sand), the cryptic pocket (green) and the cleft towards the S1 Ubiquitin binding site (blue) with shown chemical moieties. **c**, Schematic of DUB inhibition with compounds being composed of a hotspot anchor element (occupying the Ubiquitin Leu73 pocket) and one or two specificity extensions (occupying the shown other USP DUB structural elements). Compounds featuring the respective extensions are named together with their cognate DUBs.^44,46,48,53-^ 56 See Supplementary Fig. 8 for structural superpositions focused on the Leu73 binding pocket. **d**, Chemical structures of specific DUB inhibitors. Chemical motifs occupying the distinct binding sites are colored according to panel c. The hotspot anchor motifs shared by all are highlighted.

Distinctly, inhibition of USP30 by benzenesulfonamides is facilitated by a novel, inhibitor-induced conformation of the switching loop. This sets the here identified binding mode apart from all other DUB:ligand complexes. While inhibitor-bound states of UCHL1^58,59^ appear as hybrids of apo-and Ubiquitin-bound states, the novel NK036-engaged conformation prevents Ubiquitin engagement in addition to facilitating compound binding. This mechanism is also distinct from the displacement of a loop in USP1 by ML323.^52^ Our study highlights that conformational changes of the DUB need to be considered when assessing modes of inhibition and is thus complementing a recent classification of DUB inhibitor binding sites.^41^

It is striking how the hydrophobic pocket for the Ubiquitin Leu73 sidechain has emerged as a central ligandability hotspot across DUB families and how chemically diverse inhibitors are extending from there into different areas of the catalytic domains to achieve selective DUB inhibition (**Fig. 5h, Supplementary Fig. 7d**): Covalent USP7, PLpro and UCHL1 inhibitors extend towards the catalytic cysteines to bind these with electrophilic handles, whereas non-covalent USP7, USP14 and PLpro inhibitors extend into the opposite direction towards the S1 Ubiquitin binding site and the blocking loops. USP28 inhibitors extend into a pocket underneath the α5 helix, while NK036 extends into a cryptic pocket formed by the switching loop adjacent to α5.

Based on this analysis, we propose a conceptual framework for specific USP deubiquitinase inhibition which is centered on the engagement of a common ligandability hotspot in the Leu73-Ubiquitin binding site paired with diverse compound extensions (**Fig. 6c-d**). Importantly, the benzenesulfonamide scaffold stands out among DUB inhibitors with its tri-partite geometry. It demonstrates that the simultaneous presence of two such compound extensions is possible, which explains the large interaction area and high potency. The resulting distinct binding modes explain compound specificities and with more structures emerging may enable rational scaffold-hopping approaches to design novel DUB inhibitors.

Excitingly, this analysis also allowed the identification of highly related chemical moieties which occupy the hydrophobic Leu73 ligandability hotspots in structurally otherwise unrelated DUB inhibitors for different enzymes (compare the para-chlorophenyl groups in USP7 and USP14 inhibitors Compound 23 and IU1-206, respectively, and the 4-fluorophenyl group in NK036 as hotspot anchor elements, **Fig. 6d** and **Supplementary Fig. 8b-c**). This chemical feature thus appears to be a privileged structure for non-covalent DUB inhibition. Our analysis and the discovered geometry thus establish a roadmap towards specific inhibitors also for other DUBs, contributing to an emerging logic of DUB inhibition akin more established inhibitor classifications for e.g. kinases. The shared engagement of the Leu73 binding pocket in DUBs reflects the central role of Leu73 recognition for DUB activity, which is evident also from the DUB-resistant L73P mutation in ubiquitin.^62^

We here also introduce a generalizable chimeric engineering strategy to increase the crystallization propensity of USP catalytic domains. This approach expands previously explored strategies for DUB crystallization enhancement including insertion deletion for USP4, USP25, USP28, USP30,^32,53,55,63,64^ domain swapping for USP11,^65^ and surface residue mutations for USP9X, USP30^32,66^. It also synergizes with recent reports on the engineering of the Cereblon E3 ligase^67,68^ for streamlined structural characterization. We expect that our approach will form a platform to enable structure-based drug design more broadly for USP DUBs and possibly other Ubiquitin-dependent enzymes.

Collectively, our work illuminates the potential of cryptic pockets and conformational dynamics to obtain specific DUB inhibitors and advances the structure-based design of therapeutics targeting neurodegenerative diseases.

## Acknowledgments

We acknowledge beamtime at the European Synchrotron Radiation Facility (ESRF) and thank R. Gasper and P. Geue for support with crystallization and biophysics. We are grateful to the Max Planck Computing & Data Facility for the provision of computing resources. We thank all staff at the MPI Dortmund and at TU Dortmund University for excellent support. We are grateful to all members of the Gersch lab for discussions, advice, and reagents. Gifts of plasmids from D. Komander (MRC LMB Cambridge / WEHI Melbourne), G. Maertens (Imperial College London) and G. Peters (CRUK London) are gratefully acknowledged. Fig. 1a was created with BioRender. This work was funded by the Chemical Genomics Centre (AstraZeneca, Merck KGaA, Pfizer Inc., and Max Planck Society, CGCIII-352S) and the German Research Foundation through CRC1430 (DFG, 424228829). Work in the Gersch lab is further supported by an Emmy Noether grant from the German Research Foundation (DFG, GE 3110/1-1) and by the State of North Rhine-Westphalia through the ‘CANcer TARgeting Network’ (NW21-062C).

## Author contributions

NHK performed all experiments unless noted otherwise. NK synthesized chemical compounds. KG and GMK performed cellular experiments. All authors designed experiments, analyzed data, interpreted results, and generated figures. MG conceived the study, supervised research, and wrote the manuscript with input from all authors.

## Supplementary Material

Supplementary Material as a separate PDF file includes Supplementary Figures 1 to 8, Supplementary Tables 1 to 3, Protein sequences, Supplementary Methods (chemical synthesis with refs. ^33,35,69^), NMR spectra of compounds and uncropped gels and blots. Supplementary Movie 1 is supplied as a separate file.

## METHODS

### Cloning and protein expression

Chimeric USP30 constructs for bacterial expression were generated by amplifying protein-coding parts from plasmids or by incorporating sequences as overhangs into primers. The following sequences were used: Codon-optimized human USP30 catalytic domain, Addgene #110746, uniprot: Q70CQ3 with modifications described previously^32^ and in the Supplementary Material; human USP7, uniprot: Q93009; human USP14, uniprot: P54578; human USP35, uniprot: Q9P2H5; human CYLD, uniprot: Q9NQC7. All protein sequences are shown in the Supplementary Material. DNA fragments were joined through SOE-PCR and subsequently ligated into the pOPINK vector using the In-Fusion HD Cloning Kit (Takara Clonetech). A pOPINE vector containing C-terminally Flag-tagged human USP30 (1-517) was used for transient transfection as described previously.^32^ Point mutants were introduced through vector QuikChange or SOE-PCR. Constructs were confirmed by Sanger sequencing.

Bacterial expression was performed in Rosetta2(DE3)pLacI cells. An overnight culture was diluted 1:100 in 2xTY medium supplemented with appropriate antibiotics. Cultures were grown at 30°C with shaking at 180 rpm until an optical density (OD600) of 0.8-1.0 was reached. After cooling to 18°C, protein expression was induced by addition of isopropyl β-D-thiogalactopyranoside (IPTG) to a final concentration of 0.5 mM and cells were kept shaking at 18°C for 18 h. Harvested bacterial cells were stored at -80°C.

### Protein purification

Cells were resuspended in lysis buffer (50 mM sodium phosphate pH 8.0, 300 mM NaCl, 20 mM imidazole, 4 mM β-mercaptoethanol) supplemented with DNAseI and lysozyme. The cell suspension was lysed by sonication (55% amplitude, 10 s on/10 s off) on ice, cleared by centrifugation at 33,500xg for 30 min at 4°C and filtered through a 0.45 μm filter.

All purification steps were performed on Äkta Pure systems (GE Healthcare) at 4°C. For affinity chromatography, lysate was loaded onto a pre-equilibrated 5 mL HisTrap fast flow column (GE Healthcare), washed with 20 column volumes (CV) of lysis buffer and then eluted with a linear gradient of 3.5 CV into elution buffer (50 mM sodium phosphate pH 8.0, 300 mM NaCl, 500 mM imidazole, 4 mM β-mercaptoethanol). Protein-containing fractions were pooled, supplemented with His6-tagged 3C protease and dialyzed against 25 mM Tris pH 8.5, 100 mM NaCl, 4 mM DTT overnight at 4°C. The dialyzed sample was diluted two-fold with 20 mM Tris pH 8.5, filtered through a 0.45 µm filter, and directly loaded onto a 6 mL Resource Q column, equilibrated with 25 mM Tris pH 8.5, 50 mM NaCl, 4 mM DTT, for anion exchange chromatography. Elution was achieved with a gradient into high salt buffer (25 mM Tris pH 8.5, 500 mM NaCl, 4 mM DTT). Further purification was carried out on a HiLoad 16/600 Superdex 75 pg column in 20 mM Tris pH 8.0, 100 mM NaCl, 4 mM DTT for samples used for crystallography. Purity of peak fractions was assessed by SDS-PAGE before pooling, concentrating at 4°C/3200xg in spin concentrators (10 kDa MWCO, Amicon) and flash freezing in liquid nitrogen. Protein concentrations were determined on a Nanodrop (Thermo Fisher).

### Construct design and modelling

Protein construct design was guided by the annotation of USP domain boxes^39^ and available crystal structures of human USP DUBs including those of USP30^32,38^, USP7^42,45-47^, USP14^43,44^, USP35^51^ and CYLD^49^ (**Supplementary Table 1**). Chimeric USP30 protein sequences were based on the sequence used for PDB entry 5OHK.^32^ Structures of chimeric sequences were predicted with AlphaFold2^70^ and analyzed with PyMOL. Proteins were then experimentally prepared and biochemically characterized in an iterative process.

### Crystallization

Protein samples for crystallization were prepared by mixing USP30 (30 mg/mL in gel filtration buffer) with 1.4 equivalents of NK036 which was added in two steps from a 100 mM DMSO stock. Following incubation for 30 min at room temperature, the sample was passed through a Proteus mini clarification spin column (Protein Ark) and used directly for crystallization trials. Screening plates were set up by a mosquito HTS robot (TTP Labtech) in 96 well sitting-drop vapor diffusion plates in MRC format (Molecular Dimensions) and incubated at 20°C. Drop ratios of 200 nL + 200 nL and 500 nL + 500 nL were used for coarse screening and fine screening, respectively.

Initial crystals of USP30 construct ch3 in complex with NK036 were found in 0.1 M Bicine pH 9.0, 5% (w/v) PEG 20,000, 1% (v/v) dioxane. Moderately sized cylindrical crystals (60 x 16 x 16 µm^3^) were obtained through fine screening in 88 mM NaOH, 100 mM Bicine, 11% (w/v) PEG 20,000, 1% (v/v) dioxane.

Crystal size and diffraction quality were further improved through the Hampton additive screen on the above-mentioned condition. Larger cylindrical crystals (130 x 25 x 25 µm^3^) of diffraction quality were obtained in 76 mM NaOH, 100 mM Bicine, 10.2% (w/v) PEG 20,000, 1% (v/v) dioxane, 10 mM L-proline (Crystal 1) and 85 mM NaOH, 100 mM Bicine, 10.2% (w/v) PEG 20,000, 1% (v/v) dioxane, 10 mM L-proline (Crystal 2) and 88 mM NaOH, 100 mM Bicine, 11% (w/v) PEG 20,000, 1% (v/v) dioxane, 10 mM sarcosine (Crystal 3). Crystals were soaked in mother liquor supplemented with 25% (v/v) glycerol (Crystals 1 and 2) or ethylene glycol (Crystal 3) before vitrification in liquid nitrogen.

The covalent Ubiquitin complex was obtained by mixing USP30 after ResQ purification with 1.2 equivalents of Ub-PA. Following incubation at room temperature for 2 hours, USP30∼Ub-PA was purified on a HiLoad 16/600 Superdex 75 pg column as described above and concentrated to 12 mg/mL. Diffraction-quality crystals were obtained from coarse screening plates in 0.56 M sodium citrate pH 7.0. Crystals were soaked in mother liquor supplemented with 25% (v/v) ethylene glycol before vitrification in liquid nitrogen.

### Data collection, structure solution and refinement

Diffraction data were collected at 100 K at the European Synchrotron Radiation Facility (ESRF) on beamline ID30A-3. Datasets from different crystals of the USP30-NK036 complex were separately integrated with DIALS^71^. Multi-crystal data averaging was employed as implemented in CCP4 blend^72^ to improve resolution, lower *B*-factors and ultimately enhance features in the electron density. Multiple datasets and combinations were explored, resulting in the chosen data averaged from crystals 1-3 as the best cluster. The final combined data was then corrected for anisotropy by the Staraniso webserver^73^. **Table 1** displays data collection and refinement statistics, and data for anisotropically scaled individual datasets are provided in **Supplementary Table 3** for comparison. The final structure was solved to a resolution of 2.75 Å from the averaged data by molecular replacement using MR Phaser^74^ and a search model based on the Alphafold-derived structure model of the chimera. The final model was obtained by multiple cycles of structure building in Coot^75^ and refinement with Phenix Refine^76^. The compound geometry was optimized through ORCA^77^ at the B3LYP/def2-SVP level of theory. Geometry restraints were generated by ELBOW^78^.

Diffraction data leading to the USP30∼Ub-PA structure were integrated with DIALS. The final structure was obtained by molecular replacement with MR Phaser and refinement with Phenix Refine as described above. Final data collection and refinement statistics are shown in **Table 1**.

### Ub-PA labeling assay

Purified proteins were diluted in 20 mM Tris pH 8.0, 300 mM NaCl, 2 mM DTT, 5% glycerol. Protein and probe were combined at 2.5 μM and 10 μM final concentrations, respectively, and reacted for 1 h at room temperature. For the in vitro compound-probe competition assay, 1 μM protein was pre-incubated with 8 μM compound at room temperature for 2 h. Following compound incubation, 4 μM of probe was added and reacted for 1 h at 37°C. Probe binding was assessed by SDS-PAGE and Coomassie staining.

### Ub-RhoG cleavage assay

Enzyme kinetics were performed in black, low-volume, non-binding surface 384 well plates. For activity assays, 2x Ub-RhoG substrate (final concentration: 50 nM) and 2x enzyme (final concentration: 0.25 nM – 32 nM) were prepared in 20 mM HEPES pH 8.0, 50 mM NaCl, 5 mM DTT, 0.1 mg/mL BSA. 10 µL of 2x enzyme solution was added in triplicates to the plate. The reaction was started by addition of 10 µL of 2x substrate solution. For enzyme inhibition assays, 2x substrate (final concentration: 100 nM), 4x enzyme (final concentration: 0.1 nM) and 4x inhibitor (final concentration: 1.6 pM – 1.6 μM) were prepared in 50 mM Tris-HCl pH 8.0, 0.05 mg/mL BSA, 4 mM DTT, 0.01% Tween 20. 4x inhibitor solution was added to 4x enzyme solution to create a 2x enzyme-inhibitor mixture. 10 µL of this 2x enzyme-inhibitor mixture was then added to the plate in triplicates and incubated at room temperature for 90 min. The reaction was started by addition of 10 µL of 2x Ub-RhoG substrate. Substrate cleavage was monitored by measurement of cleaved RhoG fluorescence (excitation = 492 nm, emission = 525 nm) every 30 s for 1 h at 25 °C on a TECAN Spark plate reader.

### Thermal shift assay

The thermal stability of USP30 constructs and effects of inhibitor/Ub-PA binding were determined by thermal shift assays. Inhibitor, Ub-PA and Sypro orange dye (Sigma, stock: 5000x in DMSO) solutions were prepared in 1x PBS, 4 mM DTT. Inhibitor/Ub-PA were pre-diluted to 100 µM and Sypro orange dye was pre-diluted to 50x (from the commercial stock). Mixtures of protein, inhibitor/Ub-PA and Sypro were prepared having inhibitor/Ub-PA at a final concentration of 20 μM, Sypro at a final concentration of 4x and protein at a final concentration of 2 μM for c1, 4 μM for ch1, ch4 and c2 and 3 μM for ch2 and ch3. Samples were added in triplicates to a white 96-well PCR plate (Bio-Rad). Fluorescence intensity was monitored (excitation = 450-490 nm, emission = 560-580 nm) at a 20-90°C gradient (increment: 0.3 °C, hold for 5 s before read) on a Real-time PCR system (Bio-Rad).

### Cell culture

HEK293 cells (ACC 305) were obtained from the Leibniz Institute DSMZ German Collection of Microorganisms and Cell Cultures GmbH. Cells were cultured in Dulbecco’s modified Eagle’s medium (DMEM) with high glucose supplemented with 10% FBS and penicillin/streptomycin. Cells were grown at 37°C in a humidified atmosphere with 5% (v/v) CO2. Cells were tested to be free of *Mycoplasma* contamination.

### Cellular Ub-probe competition

Cells were seeded at 8 × 10^5^ cells/well and grown for 48 h. Compounds (2 µM for overexpressed USP30, 8 µM for endogenous USP30) or DMSO were added for 2 h in freshly supplied medium. The medium was aspirated, and cells washed once with ice-cold PBS. Cells were collected by scraping in lysis buffer (150 mM NaCl, 250 mM sucrose, 50 mM Tris pH 8.0, 5 mM TCEP) supplemented with EDTA-free complete protease inhibitor cocktail and benzonase. Suspended cells were homogenized by 12 bursts of a microtip probe sonicator (Branson Sonifier 450) at 10% amplitude with pulses of 1 s on ice. Remaining cell debris was separated by centrifugation (20817x*g*, 5 min, 4 °C). HA-Ub-PA probe was added to the cleared lysate (approximately 2-4 mg/mL) at a final concentration of 10 μM where indicated. After incubation for 10 min at room temperature, reactions were quenched by the addition of 4x reducing LDS sample buffer. Samples were resolved by SDS-PAGE and further analyzed via western blot.

### Western Blotting

Using a Trans-Blot Turbo system (Bio-Rad, 1.0 A, 25 V, 30 min), proteins were blotted onto a nitrocellulose membrane. Membranes were saturated by incubating in 5% (w/v) nonfat milk in PBS-T. Primary antibodies (anti-Flag, 1:1000, Sigma, F3165; anti-USP30, 1:500, Sigma, HPA016952; anti-Vinculin, 1:10000, Sigma, V9131) were allowed to bind overnight. Appropriate horseradish peroxidase-coupled secondary antibodies (anti-mouse, 1:5000, Sigma, NXA931; anti-rabbit, 1:5000, Sigma, GENA934) were applied and chemiluminescence was generated using Clarity Western ECL substrate (Bio-Rad). For detection of endogenous USP30 levels, the Clarity Western ECL substrate was supplemented with 10% Clarity Max Western ECL substrate (Bio-Rad). Images were recorded on a ChemiDoc MP Imaging system (Bio-Rad).

### Transfection

HEK293 cells (8 × 10^5^ cells/well) were seeded in six-well plates and cultivated for 24 h. PEI transfecting reagent (Polysciences) was preincubated for 10 min with the vectors in 200 μL OPTI-MEM medium (2 µg DNA per well). Cells were transfected and incubated for 48 h. Following the treatment with either compounds or DMSO in fresh media for an additional 2 h, cells were processed as described above.

### Chemical synthesis

Procedures for the synthesis of inhibitors Compound 39 and NK036 as well as compound characterization data are reported in the Supplementary Material. The synthesis was based on previously described methods^33^ for the synthesis of Compound 39, with adjustments based on the synthesis of a related compound.^69^ A different synthesis route of the title compound NK036 (termed I-137) was published elsewhere.^35^

### Data availability

Coordinates and structure factors for the USP30^ch3^ + NK036 and USP30^ch3^∼Ub-PA crystal structures were deposited with the protein databank (PDB) under accession codes 9F19 and 9F6G, respectively.

## REFERENCES

1. Bandres-Ciga, S., Diez-Fairen, M., Kim, J.J. & Singleton, A.B. Genetics of Parkinson’s disease: An introspection of its journey towards precision medicine. Neurobiol Dis 137, 104782 (2020).

2. Pickles, S., Vigie, P. & Youle, R.J. Mitophagy and Quality Control Mechanisms in Mitochondrial Maintenance. Curr Biol 28, R170–R185 (2018).

3. Jin, S.M., Lazarou, M., Wang, C., Kane, L.A., Narendra, D.P. & Youle, R.J. Mitochondrial membrane potential regulates PINK1 import and proteolytic destabilization by PARL. J Cell Biol 191, 933–42 (2010).

4. Narendra, D., Tanaka, A., Suen, D.F. & Youle, R.J. Parkin-induced mitophagy in the pathogenesis of Parkinson disease. Autophagy 5, 706–8 (2009).

5. Narendra, D.P., Jin, S.M., Tanaka, A., Suen, D.F., Gautier, C.A., Shen, J., Cookson, M.R. & Youle, R.J. PINK1 is selectively stabilized on impaired mitochondria to activate Parkin. PLoS Biol 8, e1000298 (2010).

6. Imberechts, D., Kinnart, I., Wauters, F., Terbeek, J., Manders, L., Wierda, K., Eggermont, K., Madeiro, R.F., Sue, C., Verfaillie, C. & Vandenberghe, W. DJ-1 is an essential downstream mediator in PINK1/parkin-dependent mitophagy. Brain 145, 4368–4384 (2022).

7. Borsche, M., Pereira, S.L., Klein, C. & Grunewald, A. Mitochondria and Parkinson’s Disease: Clinical, Molecular, and Translational Aspects. J Parkinsons Dis 11, 45–60 (2021).

8. Dikic, I. & Schulman, B.A. An expanded lexicon for the ubiquitin code. Nat Rev Mol Cell Biol, 1–15 (2022).

9. Harper, J.W., Ordureau, A. & Heo, J.M. Building and decoding ubiquitin chains for mitophagy. Nat Rev Mol Cell Biol 19, 93–108 (2018).

10. Nakamura, N. & Hirose, S. Regulation of mitochondrial morphology by USP30, a deubiquitinating enzyme present in the mitochondrial outer membrane. Mol Biol Cell 19, 1903–11 (2008).

11. Bingol, B., Tea, J.S., Phu, L., Reichelt, M., Bakalarski, C.E., Song, Q., Foreman, O., Kirkpatrick, D.S. & Sheng, M. The mitochondrial deubiquitinase USP30 opposes parkin-mediated mitophagy. Nature 510, 370–5 (2014).

12. Cunningham, C.N., Baughman, J.M., Phu, L., Tea, J.S., Yu, C., Coons, M., Kirkpatrick, D.S., Bingol, B. & Corn, J.E. USP30 and parkin homeostatically regulate atypical ubiquitin chains on mitochondria. Nat Cell Biol 17, 160–9 (2015).

13. Ordureau, A., Paulo, J.A., Zhang, J., An, H., Swatek, K.N., Cannon, J.R., Wan, Q., Komander, D. & Harper, J.W. Global Landscape and Dynamics of Parkin and USP30-Dependent Ubiquitylomes in iNeurons during Mitophagic Signaling. Mol Cell 77, 1124–1142 (2020).

14. Wang, F., Gao, Y., Zhou, L., Chen, J., Xie, Z., Ye, Z. & Wang, Y. USP30: Structure, Emerging Physiological Role, and Target Inhibition. Front Pharmacol 13, 851654 (2022).

15. Wang, Y., Serricchio, M., Jauregui, M., Shanbhag, R., Stoltz, T., Di Paolo, C.T., Kim, P.K. & McQuibban, G.A. Deubiquitinating enzymes regulate PARK2-mediated mitophagy. Autophagy 11, 595–606 (2015).

16. Ordureau, A., Sarraf, S.A., Duda, D.M., Heo, J.M., Jedrychowski, M.P., Sviderskiy, V.O., Olszewski, J.L., Koerber, J.T., Xie, T., Beausoleil, S.A., Wells, J.A., Gygi, S.P., Schulman, B.A. & Harper, J.W. Quantitative proteomics reveal a feedforward mechanism for mitochondrial PARKIN translocation and ubiquitin chain synthesis. Mol Cell 56, 360–375 (2014).

17. Marcassa, E., Kallinos, A., Jardine, J., Rusilowicz-Jones, E.V., Martinez, A., Kuehl, S., Islinger, M., Clague, M.J. & Urbe, S. Dual role of USP30 in controlling basal pexophagy and mitophagy. EMBO Rep 19, e45595 (2018).

18. Sanchez-Martinez, A., Martinez, A. & Whitworth, A.J. FBXO7/ntc and USP30 antagonistically set the ubiquitination threshold for basal mitophagy and provide a target for Pink1 phosphorylation in vivo. PLoS Biol 21, e3002244 (2023).

19. Phu, L., Rose, C.M., Tea, J.S., Wall, C.E., Verschueren, E., Cheung, T.K., Kirkpatrick, D.S. & Bingol, B. Dynamic Regulation of Mitochondrial Import by the Ubiquitin System. Mol Cell 77, 1107–1123 (2020).

20. Yue, W., Chen, Z., Liu, H., Yan, C., Chen, M., Feng, D., Yan, C., Wu, H., Du, L., Wang, Y., Liu, J., Huang, X., Xia, L., Liu, L., Wang, X., Jin, H., Wang, J., Song, Z., Hao, X. & Chen, Q. A small natural molecule promotes mitochondrial fusion through inhibition of the deubiquitinase USP30. Cell Res 24, 482–96 (2014).

21. Lisci, M., Barton, P.R., Randzavola, L.O., Ma, C.Y., Marchingo, J.M., Cantrell, D.A., Paupe, V., Prudent, J., Stinchcombe, J.C. & Griffiths, G.M. Mitochondrial translation is required for sustained killing by cytotoxic T cells. Science 374, eabe9977 (2021).

22. Liang, J.R., Martinez, A., Lane, J.D., Mayor, U., Clague, M.J. & Urbe, S. USP30 deubiquitylates mitochondrial Parkin substrates and restricts apoptotic cell death. EMBO Rep 16, 618–27 (2015).

23. Riccio, V., Demers, N., Hua, R., Vissa, M., Cheng, D.T., Strilchuk, A.W., Wang, Y., McQuibban, G.A. & Kim, P.K. Deubiquitinating enzyme USP30 maintains basal peroxisome abundance by regulating pexophagy. J Cell Biol 218, 798–807 (2019).

24. Fang, T.Z., Sun, Y., Pearce, A.C., Eleuteri, S., Kemp, M., Luckhurst, C.A., Williams, R., Mills, R., Almond, S., Burzynski, L., Markus, N.M., Lelliott, C.J., Karp, N.A., Adams, D.J., Jackson, S.P., Zhao, J.F., Ganley, I.G., Thompson, P.W., Balmus, G. & Simon, D.K. Knockout or inhibition of USP30 protects dopaminergic neurons in a Parkinson’s disease mouse model. Nat Commun 14, 7295 (2023).

25. Okarmus, J., Agergaard, J.B., Stummann, T.C., Haukedal, H., Ambjorn, M., Freude, K.K., Fog, K. & Meyer, M. USP30 inhibition induces mitophagy and reduces oxidative stress in parkin-deficient human neurons. Cell Death Dis 15, 52 (2024).

26. Miller, S. & Muqit, M.M.K. Therapeutic approaches to enhance PINK1/Parkin mediated mitophagy for the treatment of Parkinson’s disease. Neurosci Lett 705, 7–13 (2019).

27. Rusilowicz-Jones, E.V., Barone, F.G., Lopes, F.M., Stephen, E., Mortiboys, H., Urbe, S. & Clague, M.J. Benchmarking a highly selective USP30 inhibitor for enhancement of mitophagy and pexophagy. Life Sci Alliance 5, e202101287 (2022).

28. Rusilowicz-Jones, E.V., Jardine, J., Kallinos, A., Pinto-Fernandez, A., Guenther, F., Giurrandino, M., Barone, F.G., McCarron, K., Burke, C.J., Murad, A., Martinez, A., Marcassa, E., Gersch, M., Buckmelter, A.J., Kayser-Bricker, K.J., Lamoliatte, F., Gajbhiye, A., Davis, S., Scott, H.C., Murphy, E., England, K., Mortiboys, H., Komander, D., Trost, M., Kessler, B.M., Ioannidis, S., Ahlijanian, M.K., Urbe, S. & Clague, M.J. USP30 sets a trigger threshold for PINK1-PARKIN amplification of mitochondrial ubiquitylation. Life Sci Alliance 3, e202000768 (2020).

29. Agarwal, A., Dong, Z., Harris, R., Murray, P., Parikh, S.M., Rosner, M.H., Kellum, J.A., Ronco, C. & Acute Dialysis Quality Initiative, X.W.G. Cellular and Molecular Mechanisms of AKI. J Am Soc Nephrol 27, 1288–99 (2016).

30. Ferrer, S., Muratore, M.E. & Buijnsters, P. The intriguing role of USP30 inhibitors as deubiquitinating enzymes from the patent literature since 2013. Expert Opin Ther Pat 32, 523–559 (2022).

31. Mondal, M., Cao, F., Conole, D., Auner, H.W. & Tate, E.W. Discovery of potent and selective activity-based probes (ABPs) for the deubiquitinating enzyme USP30. RSC Chem Biol 5, 439–446 (2024).

32. Gersch, M., Gladkova, C., Schubert, A.F., Michel, M.A., Maslen, S. & Komander, D. Mechanism and regulation of the Lys6-selective deubiquitinase USP30. Nat Struct Mol Biol 24, 920–930 (2017).

33. Kluge, A.F., Lagu, B.R., Maiti, P., Jaleel, M., Webb, M., Malhotra, J., Mallat, A., Srinivas, P.A. & Thompson, J.E. Novel highly selective inhibitors of ubiquitin specific protease 30 (USP30) accelerate mitophagy. Bioorg Med Chem Lett 28, 2655–2659 (2018).

34. O’Brien, D.P., Jones, H.B.L., Guenther, F., Murphy, E.J., England, K.S., Vendrell, I., Anderson, M., Brennan, P.E., Davis, J.B., Pinto-Fernandez, A., Turnbull, A.P. & Kessler, B.M. Structural Premise of Selective Deubiquitinase USP30 Inhibition by Small-Molecule Benzosulfonamides. Mol Cell Proteomics 22, 100609 (2023).

35. Romero, D. L.; Johnson, M., Garrett Lee, A., David Behrouz, B. & Fritzen Jr., E., Lawrence. USP30 Inhibitors and uses thereof. WO/2021/050992 (2021).

36. Luo, H., Krigman, J., Zhang, R., Yang, M. & Sun, N. Pharmacological inhibition of USP30 activates tissue-specific mitophagy. Acta Physiol (Oxf) 232, e13666 (2021).

37. Barone, F.G., Urbe, S. & Clague, M.J. Segregation of pathways leading to pexophagy. Life Sci Alliance 6, e202201825 (2023).

38. Sato, Y., Okatsu, K., Saeki, Y., Yamano, K., Matsuda, N., Kaiho, A., Yamagata, A., Goto-Ito, S., Ishikawa, M., Hashimoto, Y., Tanaka, K. & Fukai, S. Structural basis for specific cleavage of Lys6-linked polyubiquitin chains by USP30. Nat Struct Mol Biol 24, 911–919 (2017).

39. Ye, Y., Scheel, H., Hofmann, K. & Komander, D. Dissection of USP catalytic domains reveals five common insertion points. Mol BioSyst 5, 1797–808 (2009).

40. Clague, M.J., Urbe, S. & Komander, D. Breaking the chains: deubiquitylating enzyme specificity begets function. Nat Rev Mol Cell Biol 20, 338–352 (2019).

41. Lange, S.M., Armstrong, L.A. & Kulathu, Y. Deubiquitinases: From mechanisms to their inhibition by small molecules. Mol Cell 82, 15–29 (2022).

42. Hu, M., Li, P., Li, M., Li, W., Yao, T., Wu, J.W., Gu, W., Cohen, R.E. & Shi, Y. Crystal structure of a UBP-family deubiquitinating enzyme in isolation and in complex with ubiquitin aldehyde. Cell 111, 1041–54 (2002).

43. Hu, M., Li, P., Song, L., Jeffrey, P.D., Chenova, T.A., Wilkinson, K.D., Cohen, R.E. & Shi, Y. Structure and mechanisms of the proteasome-associated deubiquitinating enzyme USP14. EMBO J 24, 3747–56 (2005).

44. Wang, Y., Jiang, Y., Ding, S., Li, J., Song, N., Ren, Y., Hong, D., Wu, C., Li, B., Wang, F., He, W., Wang, J. & Mei, Z. Small molecule inhibitors reveal allosteric regulation of USP14 via steric blockade. Cell Res 28, 1186–1194 (2018).

45. Kategaya, L., Di Lello, P., Rouge, L., Pastor, R., Clark, K.R., Drummond, J., Kleinheinz, T., Lin, E., Upton, J.P., Prakash, S., Heideker, J., McCleland, M., Ritorto, M.S., Alessi, D.R., Trost, M., Bainbridge, T.W., Kwok, M.C.M., Ma, T.P., Stiffler, Z., Brasher, B., Tang, Y., Jaishankar, P., Hearn, B.R., Renslo, A.R., Arkin, M.R., Cohen, F., Yu, K., Peale, F., Gnad, F., Chang, M.T., Klijn, C., Blackwood, E., Martin, S.E., Forrest, W.F., Ernst, J.A., Ndubaku, C., Wang, X., Beresini, M.H., Tsui, V., Schwerdtfeger, C., Blake, R.A., Murray, J., Maurer, T. & Wertz, I.E. USP7 small-molecule inhibitors interfere with ubiquitin binding. Nature 550, 534–538 (2017).

46. Turnbull, A.P., Ioannidis, S., Krajewski, W.W., Pinto-Fernandez, A., Heride, C., Martin, A.C.L., Tonkin, L.M., Townsend, E.C., Buker, S.M., Lancia, D.R., Caravella, J.A., Toms, A.V., Charlton, T.M., Lahdenranta, J., Wilker, E., Follows, B.C., Evans, N.J., Stead, L., Alli, C., Zarayskiy, V.V., Talbot, A.C., Buckmelter, A.J., Wang, M., McKinnon, C.L., Saab, F., McGouran, J.F., Century, H., Gersch, M., Pittman, M.S., Marshall, C.G., Raynham, T.M., Simcox, M., Stewart, L.M.D., McLoughlin, S.B., Escobedo, J.A., Bair, K.W., Dinsmore, C.J., Hammonds, T.R., Kim, S., Urbe, S., Clague, M.J., Kessler, B.M. & Komander, D. Molecular basis of USP7 inhibition by selective small-molecule inhibitors. Nature 550, 481–486 (2017).

47. Lamberto, I., Liu, X., Seo, H.S., Schauer, N.J., Iacob, R.E., Hu, W., Das, D., Mikhailova, T., Weisberg, E.L., Engen, J.R., Anderson, K.C., Chauhan, D., Dhe-Paganon, S. & Buhrlage, S.J. Structure-Guided Development of a Potent and Selective Non-covalent Active-Site Inhibitor of USP7. Cell Chem Biol 24, 1490–1500 (2017).

48. Gavory, G., O’Dowd, C.R., Helm, M.D., Flasz, J., Arkoudis, E., Dossang, A., Hughes, C., Cassidy, E., McClelland, K., Odrzywol, E., Page, N., Barker, O., Miel, H. & Harrison, T. Discovery and characterization of highly potent and selective allosteric USP7 inhibitors. Nat Chem Biol 14, 118–125 (2018).

49. Komander, D., Lord, C.J., Scheel, H., Swift, S., Hofmann, K., Ashworth, A. & Barford, D. The structure of the CYLD USP domain explains its specificity for Lys63-linked polyubiquitin and reveals a B box module. Mol Cell 29, 451–64 (2008).

50. Sato, Y., Goto, E., Shibata, Y., Kubota, Y., Yamagata, A., Goto-Ito, S., Kubota, K., Inoue, J., Takekawa, M., Tokunaga, F. & Fukai, S. Structures of CYLD USP with Met1-or Lys63-linked diubiquitin reveal mechanisms for dual specificity. Nat Struct Mol Biol 22, 222–9 (2015).

51. Leznicki, P., Natarajan, J., Bader, G., Spevak, W., Schlattl, A., Abdul Rehman, S.A., Pathak, D., Weidlich, S., Zoephel, A., Bordone, M.C., Barbosa-Morais, N.L., Boehmelt, G. & Kulathu, Y. Expansion of DUB functionality generated by alternative isoforms - USP35, a case study. J Cell Sci 131, jcs212753 (2018).

52. Rennie, M.L., Arkinson, C., Chaugule, V.K. & Walden, H. Cryo-EM reveals a mechanism of USP1 inhibition through a cryptic binding site. Sci Adv 8, eabq6353 (2022).

53. Patzke, J.V., Sauer, F., Nair, R.K., Endres, E., Proschak, E., Hernandez-Olmos, V., Sotriffer, C. & Kisker, C. Structural basis for the bi-specificity of USP25 and USP28 inhibitors. EMBO Rep (2024).

54. Leger, P.R., Hu, D.X., Biannic, B., Bui, M., Han, X., Karbarz, E., Maung, J., Okano, A., Osipov, M., Shibuya, G.M., Young, K., Higgs, C., Abraham, B., Bradford, D., Cho, C., Colas, C., Jacobson, S., Ohol, Y.M., Pookot, D., Rana, P., Sanchez, J., Shah, N., Sun, M., Wong, S., Brockstedt, D.G., Kassner, P.D., Schwarz, J.B. & Wustrow, D.J. Discovery of Potent, Selective, and Orally Bioavailable Inhibitors of USP7 with In Vivo Antitumor Activity. J Med Chem 63, 5398–5420 (2020).

55. Zhou, D., Xu, Z., Huang, Y., Wang, H., Zhu, X., Zhang, W., Song, W., Gao, T., Liu, T., Wang, M., Shi, L., Zhang, N. & Xiong, B. Structure-based discovery of potent USP28 inhibitors derived from Vismodegib. Eur J Med Chem 254, 115369 (2023).

56. Baez-Santos, Y.M., Barraza, S.J., Wilson, M.W., Agius, M.P., Mielech, A.M., Davis, N.M., Baker, S.C., Larsen, S.D. & Mesecar, A.D. X-ray structural and biological evaluation of a series of potent and highly selective inhibitors of human coronavirus papain-like proteases. J Med Chem 57, 2393–412 (2014).

57. Sanders, B.C., Pokhrel, S., Labbe, A.D., Mathews, II, Cooper, C.J., Davidson, R.B., Phillips, G., Weiss, K.L., Zhang, Q., O’Neill, H., Kaur, M., Schmidt, J.G., Reichard, W., Surendranathan, S., Parvathareddy, J., Phillips, L., Rainville, C., Sterner, D.E., Kumaran, D., Andi, B., Babnigg, G., Moriarty, N.W., Adams, P.D., Joachimiak, A., Hurst, B.L., Kumar, S., Butt, T.R., Jonsson, C.B., Ferrins, L., Wakatsuki, S., Galanie, S., Head, M.S. & Parks, J.M. Potent and selective covalent inhibition of the papain-like protease from SARS-CoV-2. Nat Commun 14, 1733 (2023).

58. Grethe, C., Schmidt, M., Kipka, G.M., O’Dea, R., Gallant, K., Janning, P. & Gersch, M. Structural basis for specific inhibition of the deubiquitinase UCHL1. Nat Commun 13, 5950 (2022).

59. Schmidt, M., Grethe, C., Recknagel, S., Kipka, G.M., Klink, N. & Gersch, M. N-Cyanopiperazines as Specific Covalent Inhibitors of the Deubiquitinating Enzyme UCHL1. Angew Chem Int Ed Engl 63, e202318849 (2024).

60. Wertz, I.E. & Wang, X. From Discovery to Bedside: Targeting the Ubiquitin System. Cell Chem Biol 26, 156–177 (2019).

61. Scientists associated with Amgen Inc and Carmot Therapeutics Inc have deposited two crystal structures of human USP30 construct 1 in complex with a custom antibody and two covalent inhibitors in the PDB (8D0A, 8D1T). While the citation box states „to be published”, we have not identified a peer-reviewed or pre-printed manuscript by the authors covering these structures. The different chemical scaffold and different binding mode make these geometries complementary to our study. However, in the absence of any validation work, compound potency or specifcity, antibody characterisation data, or interpretation, we are reluctant to draw comparisons. We look forward to discussing all structures side-by-side possibly in a revised version of this manuscript or a future study.

62. Bekes, M., Okamoto, K., Crist, S.B., Jones, M.J., Chapman, J.R., Brasher, B.B., Melandri, F.D., Ueberheide, B.M., Denchi, E.L. & Huang, T.T. DUB-resistant ubiquitin to survey ubiquitination switches in mammalian cells. Cell Rep 5, 826–38 (2013).

63. Clerici, M., Luna-Vargas, M.P., Faesen, A.C. & Sixma, T.K. The DUSP-Ubl domain of USP4 enhances its catalytic efficiency by promoting ubiquitin exchange. Nat Commun 5, 5399 (2014).

64. Gersch, M., Wagstaff, J.L., Toms, A.V., Graves, B., Freund, S.M.V. & Komander, D. Distinct USP25 and USP28 Oligomerization States Regulate Deubiquitinating Activity. Mol Cell 74, 436–451 (2019).

65. Maurer, S.K., Mayer, M.P., Ward, S.J., Boudjema, S., Halawa, M., Zhang, J., Caulton, S.G., Emsley, J. & Dreveny, I. Ubiquitin-specific protease 11 structure in complex with an engineered substrate mimetic reveals a molecular feature for deubiquitination selectivity. J Biol Chem 299, 105300 (2023).

66. Paudel, P., Zhang, Q., Leung, C., Greenberg, H.C., Guo, Y., Chern, Y.H., Dong, A., Li, Y., Vedadi, M., Zhuang, Z. & Tong, Y. Crystal structure and activity-based labeling reveal the mechanisms for linkage-specific substrate recognition by deubiquitinase USP9X. Proc Natl Acad Sci U S A 116, 7288–7297 (2019).

67. Kroupova, A., Spiteri, V.A., Furihata, H., Darren, D., Ramachandran, S., Rutter, Z.J., Chakraborti, S., Haubrich, K., Pethe, J., Gonzales, D., Wijaya, A., Rodriguez-Rios, M., Lynch, D.M., Farnaby, W., Nakasone, M.A., Zollman, D. & Ciulli, A. Design of a Cereblon construct for crystallographic and biophysical studies of protein degraders. bioRxiv, 2024.01.17.575503 (2024).

68. Bailey, H.J., Eisert, J., Vollrath, J., Leibrock, E.-M., Kondratov, I., Matviyuk, T., Tolmachova, N., Langer, J.D., Wegener, A.A., Sorrell, F.J. & Dikic, I. Engineering CRBN for rapid identification of next generation binders. bioRxiv, 2024.01.18.576231 (2024).

69. Varca, A.C., Casalena, D., Chan, W.C., Hu, B., Magin, R.S., Roberts, R.M., Liu, X., Zhu, H., Seo, H.S., Dhe-Paganon, S., Marto, J.A., Auld, D. & Buhrlage, S.J. Identification and validation of selective deubiquitinase inhibitors. Cell Chem Biol 28, 1758–1771 e13 (2021).

70. Jumper, J., Evans, R., Pritzel, A., Green, T., Figurnov, M., Ronneberger, O., Tunyasuvunakool, K., Bates, R., Zidek, A., Potapenko, A., Bridgland, A., Meyer, C., Kohl, S.A.A., Ballard, A.J., Cowie, A., Romera-Paredes, B., Nikolov, S., Jain, R., Adler, J., Back, T., Petersen, S., Reiman, D., Clancy, E., Zielinski, M., Steinegger, M., Pacholska, M., Berghammer, T., Bodenstein, S., Silver, D., Vinyals, O., Senior, A.W., Kavukcuoglu, K., Kohli, P. & Hassabis, D. Highly accurate protein structure prediction with AlphaFold. Nature 596, 583–589 (2021).

71. Winter, G., Waterman, D.G., Parkhurst, J.M., Brewster, A.S., Gildea, R.J., Gerstel, M., Fuentes-Montero, L., Vollmar, M., Michels-Clark, T., Young, I.D., Sauter, N.K. & Evans, G. DIALS: implementation and evaluation of a new integration package. Acta Crystallogr D Struct Biol 74, 85–97 (2018).

72. Foadi, J., Aller, P., Alguel, Y., Cameron, A., Axford, D., Owen, R.L., Armour, W., Waterman, D.G., Iwata, S. & Evans, G. Clustering procedures for the optimal selection of data sets from multiple crystals in macromolecular crystallography. Acta Crystallogr D Struct Biol 69, 1617–32 (2013).

73. Tickle, I.J., Flensburg, C., Keller, P., Paciorek, W., Sharff, A., Vonrhein, C. & Bricogne, G. STARANISO (staraniso.globalphasing.org/). Cambridge, United Kingdom: Global Phasing Ltd. (2016).

74. McCoy, A.J., Grosse-Kunstleve, R.W., Adams, P.D., Winn, M.D., Storoni, L.C. & Read, R.J. Phaser crystallographic software. J Appl Crystallogr 40, 658–674 (2007).

75. Emsley, P., Lohkamp, B., Scott, W.G. & Cowtan, K. Features and development of Coot. Acta Crystallogr D Struct Biol 66, 486–501 (2010).

76. Adams, P.D., Afonine, P.V., Bunkoczi, G., Chen, V.B., Echols, N., Headd, J.J., Hung, L.W., Jain, S., Kapral, G.J., Grosse Kunstleve, R.W., McCoy, A.J., Moriarty, N.W., Oeffner, R.D., Read, R.J., Richardson, D.C., Richardson, J.S., Terwilliger, T.C. & Zwart, P.H. The Phenix software for automated determination of macromolecular structures. Methods 55, 94–106 (2011).

77. Neese, F., Wennmohs, F., Becker, U. & Riplinger, C. The ORCA quantum chemistry program package. J Chem Phys 152, 224108 (2020).

78. Moriarty, N.W., Grosse-Kunstleve, R.W. & Adams, P.D. electronic Ligand Builder and Optimization Workbench (eLBOW): a tool for ligand coordinate and restraint generation. Acta Crystallogr D Struct Biol 65, 1074–80 (2009).

