## Supplementary Information for "Chimeric deubiquitinase engineering reveals structural basis for specific inhibition of USP30 and a framework for DUB ligandability"

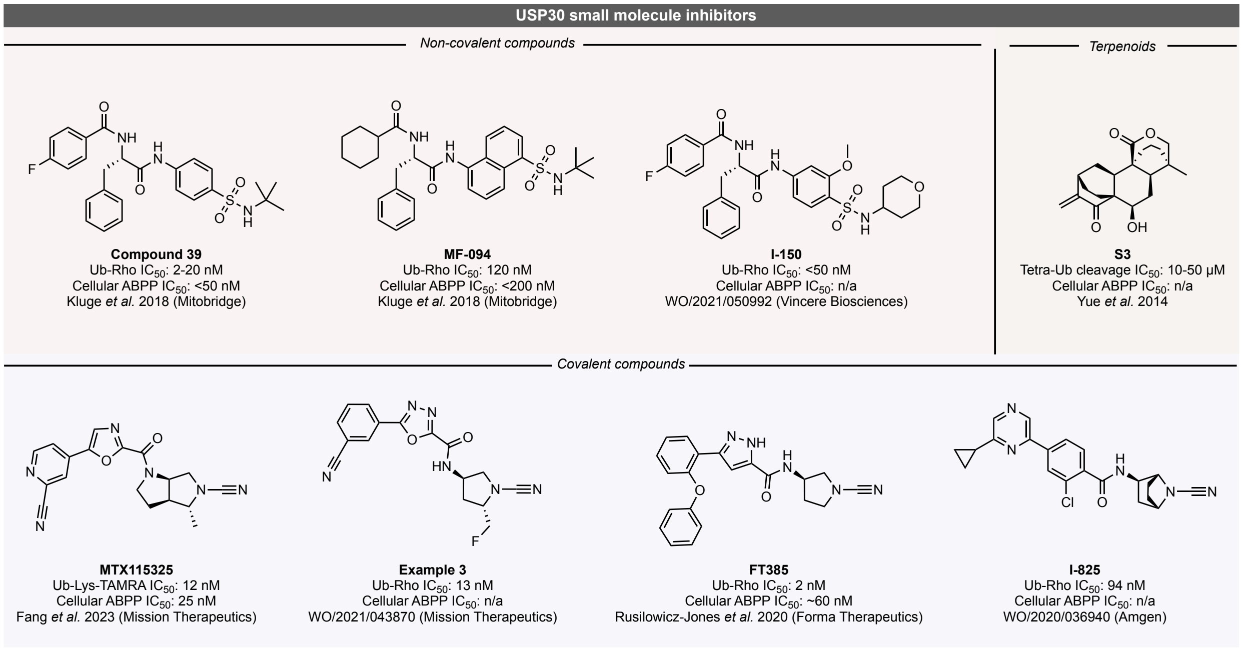

#### Supplementary Fig. 1 | Small molecule inhibitors of USP30.

Chemical structures of representative examples of small molecule USP30 inhibitors are given together with characterization data and references. Cyanamides of covalent inhibitors form isothiourea linkages with the active site cysteine of USP30. Non-covalent inhibitors belong either to chemical series which were developed from phenylalanine derivatives (with either a benzenesulfonamide or a naphthylsulfonamide) or are natural product derivatives.

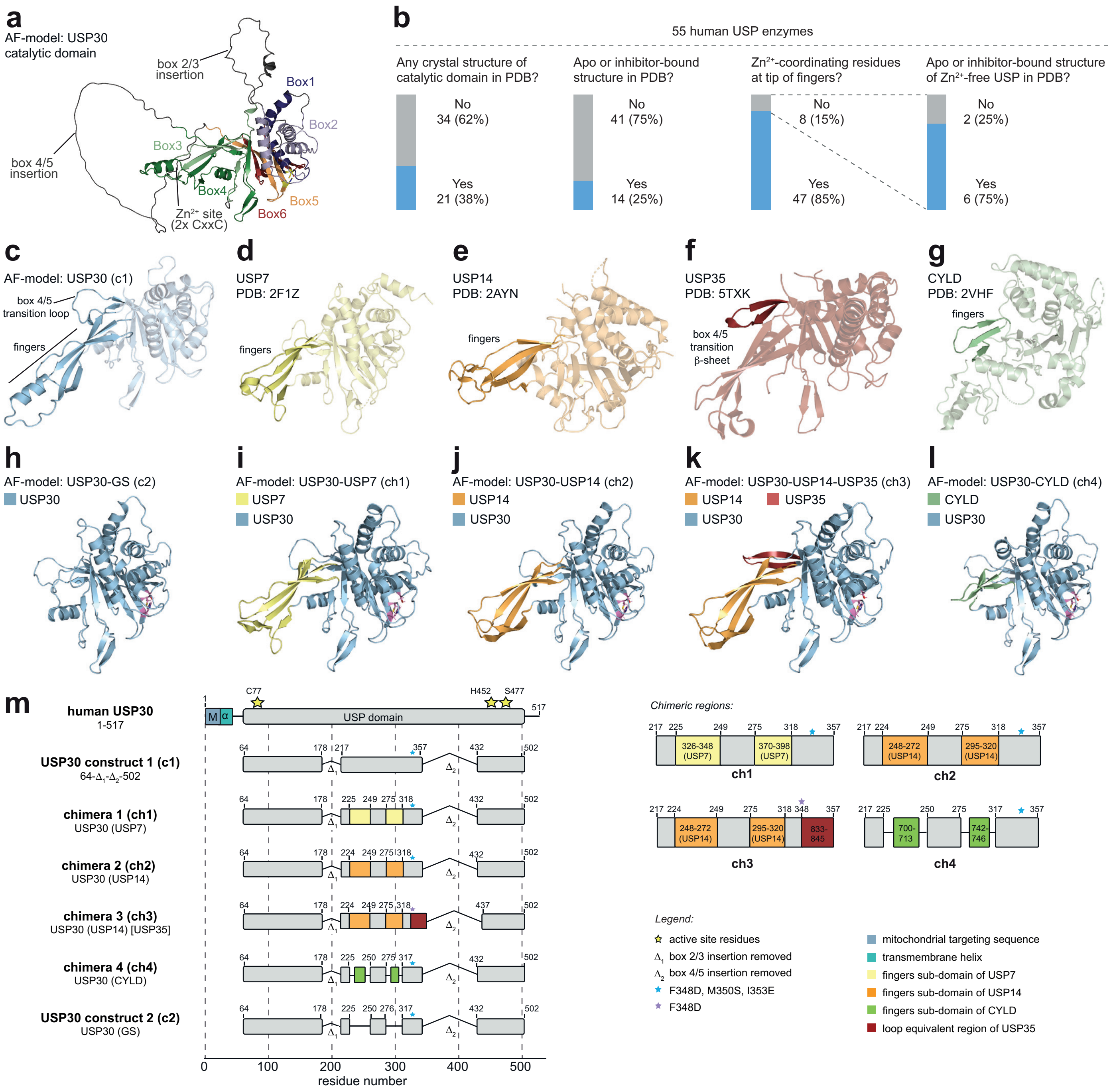

#### Supplementary Fig. 2 | Design of chimeric USP30 protein constructs.

**a,** Alphafold (AF)-model of the catalytic domain of human USP30. USP boxes and domain insertions are indicated. **b,** Statistics of human USP enzymes regarding their characterization by crystallography and regarding the presence of $Zn^{2+}$-coordinating residues in the tip of the fingers subdomain. **c,** AF-model of USP30$^{c1}$, previously optimized for structural studies and used as a starting point for this project. **d-g,** Experimental structures of catalytic domains of USP7 (d), USP14 (e), USP35 (f), and CYLD (g). Elements used for USP30 chimeric engineering are highlighted. **h-l,** AF-models of USP30 constructs explored in this study. Catalytic residues are shown in pink. **m,** Architecture of USP30 constructs with close-up view of the boundaries of the chimeric portions.

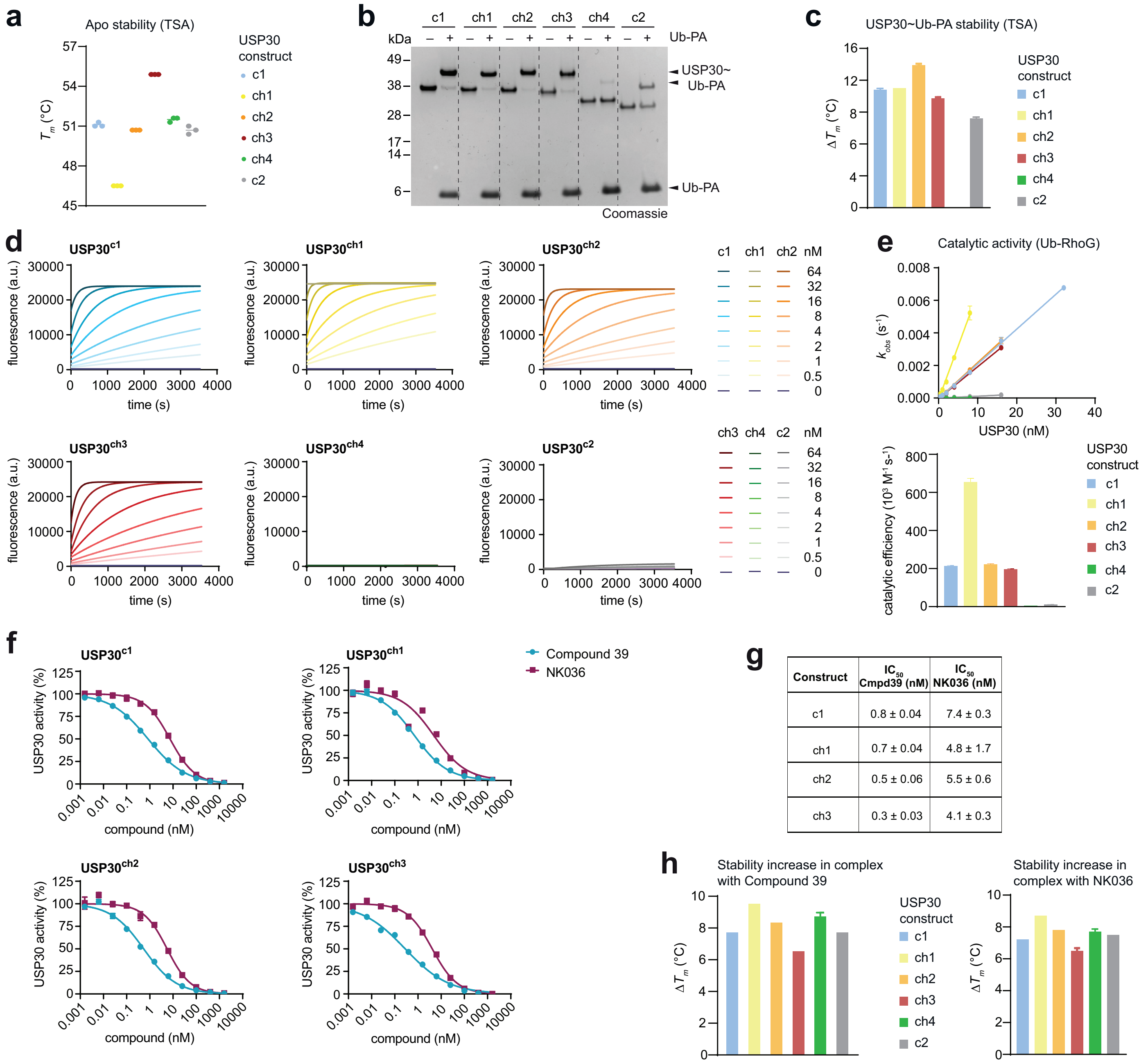

**g**

| Construct | $IC_{50}$ Cmpd39 (nM) | $IC_{50}$ NK036 (nM) |
|---|---|---|
| c1 | 0.8 ± 0.04 | 7.4 ± 0.3 |
| ch1 | 0.7 ± 0.04 | 4.8 ± 1.7 |
| ch2 | 0.5 ± 0.06 | 5.5 ± 0.6 |
| ch3 | 0.3 ± 0.03 | 4.1 ± 0.3 |

#### Supplementary Fig. 3 | Biochemical characterization of USP30 chimeras.

**a,** Protein stability assessment of USP30 constructs with thermal shift assays. **b,** Ubiquitin probe reactivity assay. Samples were analyzed by SDS-PAGE and Coomassie staining. **c,** Changes in protein stability upon binding to Ub-PA. $\Delta T_m$ was calculated as $T_m$ (Ub-PA-bound) subtracted from $T_m$ (apo protein). Mean ± s.d. (N=3). **d,** Quantification of enzyme activity. Varying concentrations of USP30 proteins were incubated with Ubiquitin-RhoG substrate and fluorescence was recorded. **e,** Observed rate constants derived from plots in d were plotted over enzyme concentrations (upper panel) to derive catalytic efficiencies (lower panel). Mean ± s.e.m. **f,** Inhibitory potencies of Compound 39 and NK036. Compounds were pre-incubated with USP30 constructs for 1.5 h, and remaining activities were determined from Ub-RhoG cleavage assays. **g,** $IC_{50}$ values of assays shown in f. **h,** Assessment of binding of Compound 39 and NK036 to indicated USP30 constructs by thermal shift assays. $\Delta T_m$ was calculated as $T_m$ of the inhibitor-bound sample subtracted from the $T_m$ of the apo protein.

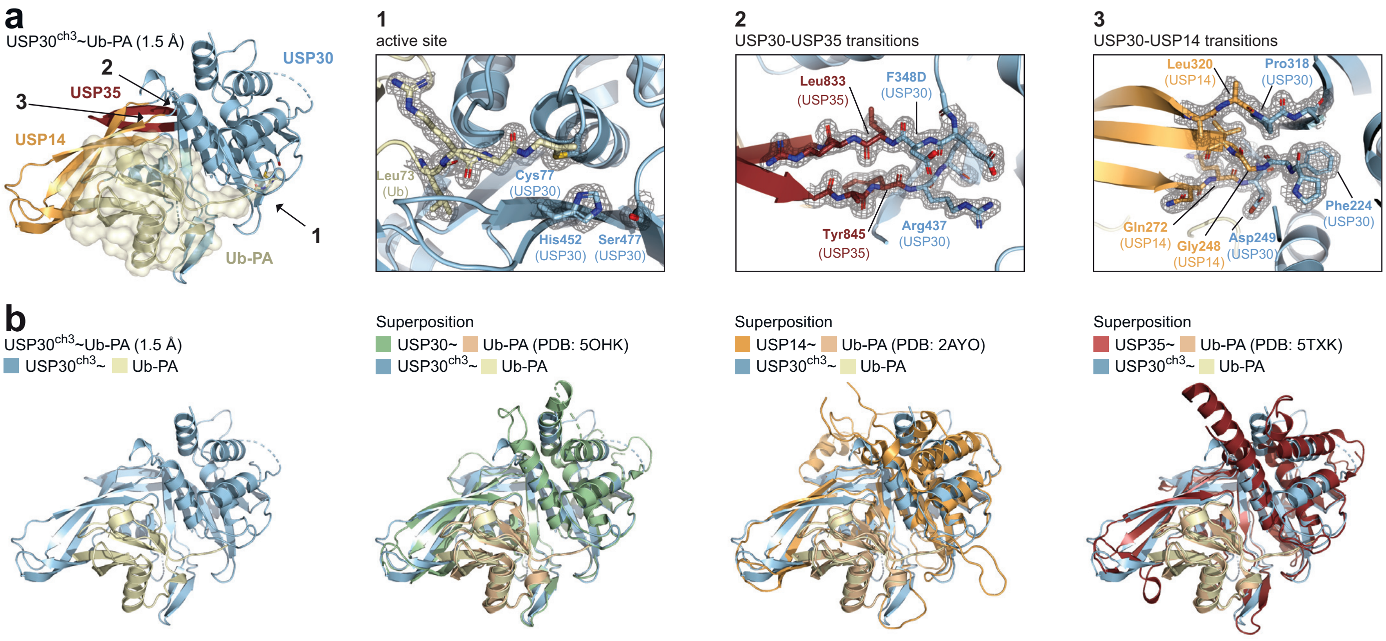

#### Supplementary Fig. 4 | Crystal structure of USP30$^{ch3}$ in complex with Ub-PA.

**a**, Cartoon representation of crystal structure of USP30$^{ch3}$~Ub-PA, with representative $2F_O$-$F_C$ electron density (contoured at 1σ) shown for indicated areas. **b**, USP30$^{ch3}$~Ub-PA structure and superpositions with previously reported Ub-PA complexes of USP30$^{c1}$, USP14 and USP35.

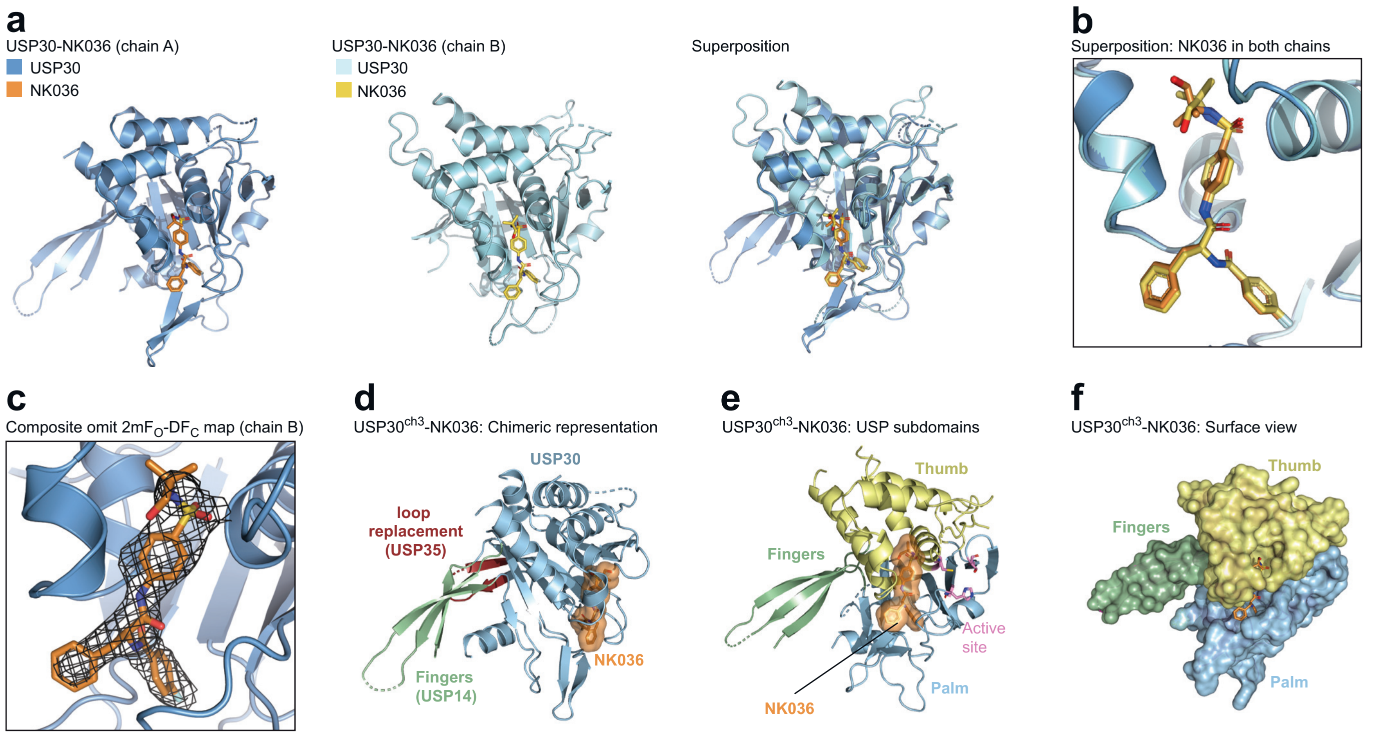

#### Supplementary Fig. 5 | Crystal structure of USP30 in complex with NK036.

**a,** Cartoon representations of the two copies within the asymmetric unit in the crystal structure of USP30$^{ch3}$-NK036. **b,** Close-up view of the superposition shown in a highlighting the ligand geometries in both chains. **c,** Composite omit electron density map of NK036 in chain B (2mF$_O$-DF$_C$, contoured at 1σ, covering all atoms of the compound). **d,** Structure as in a, with chimeric elements shown in different colors. Of note, no chimeric residue is near the compound binding site. **e,** Cartoon representation of crystal structure of USP30$^{ch3}$-NK036, highlighting different USP subdomains. The compound is shown under an orange surface, active site residues are shown in pink. **f,** Structure as in panel e with surface representation of USP30 highlighting different USP subdomains.

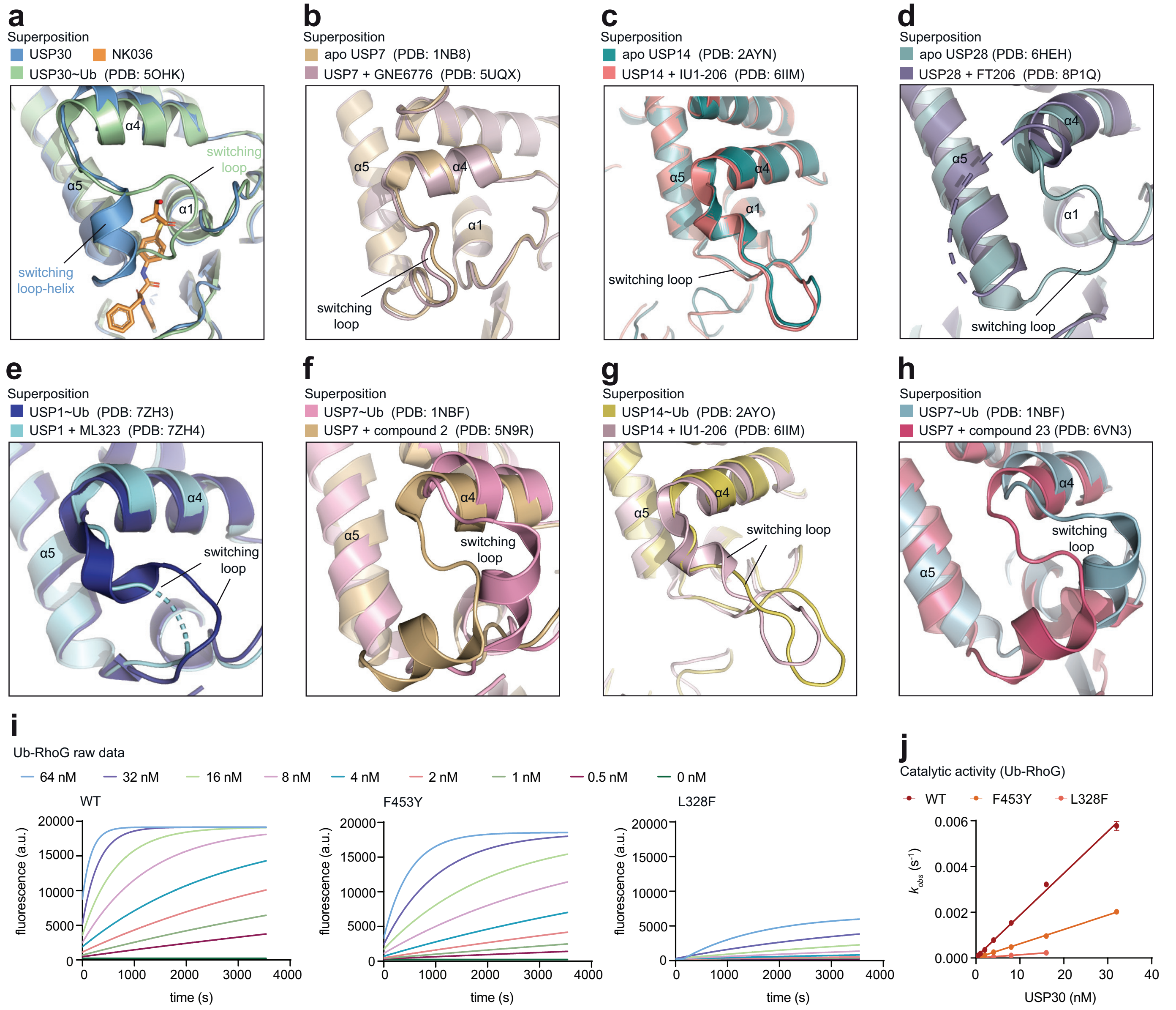

#### Supplementary Fig. 6 | Molecular basis of NK036 potency and specificity towards USP30.

**a-h,** Structural superpositions showing the switching loop positions in different USP DUBs, comparing compound-bound to ubiquitin-bound or apo states. **i-j,** Quantification of USP30 enzyme activity for WT and indicated mutant USP30, as described in Supplementary Fig. 3d-e.

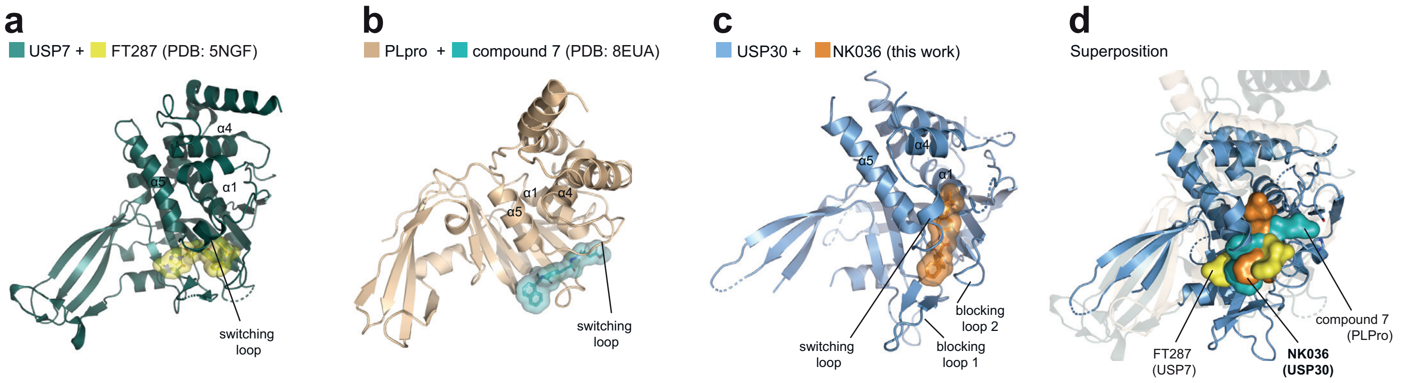

**Supplementary Fig. 7 | Comparison of the binding mode of NK036 to covalent USP DUB inhibitors.**

**a-c,** Cartoon representations of the crystal structures of USP7, SARS-CoV2-PLPro and USP30 in complex with respective inhibitors. Compounds are shown as surfaces. **d,** Superposition of the structure of USP30+NK036 on other structures shown in panels a-c. Compounds are shown as surfaces and are labeled. All protein cartoons except USP30 are semitransparent.

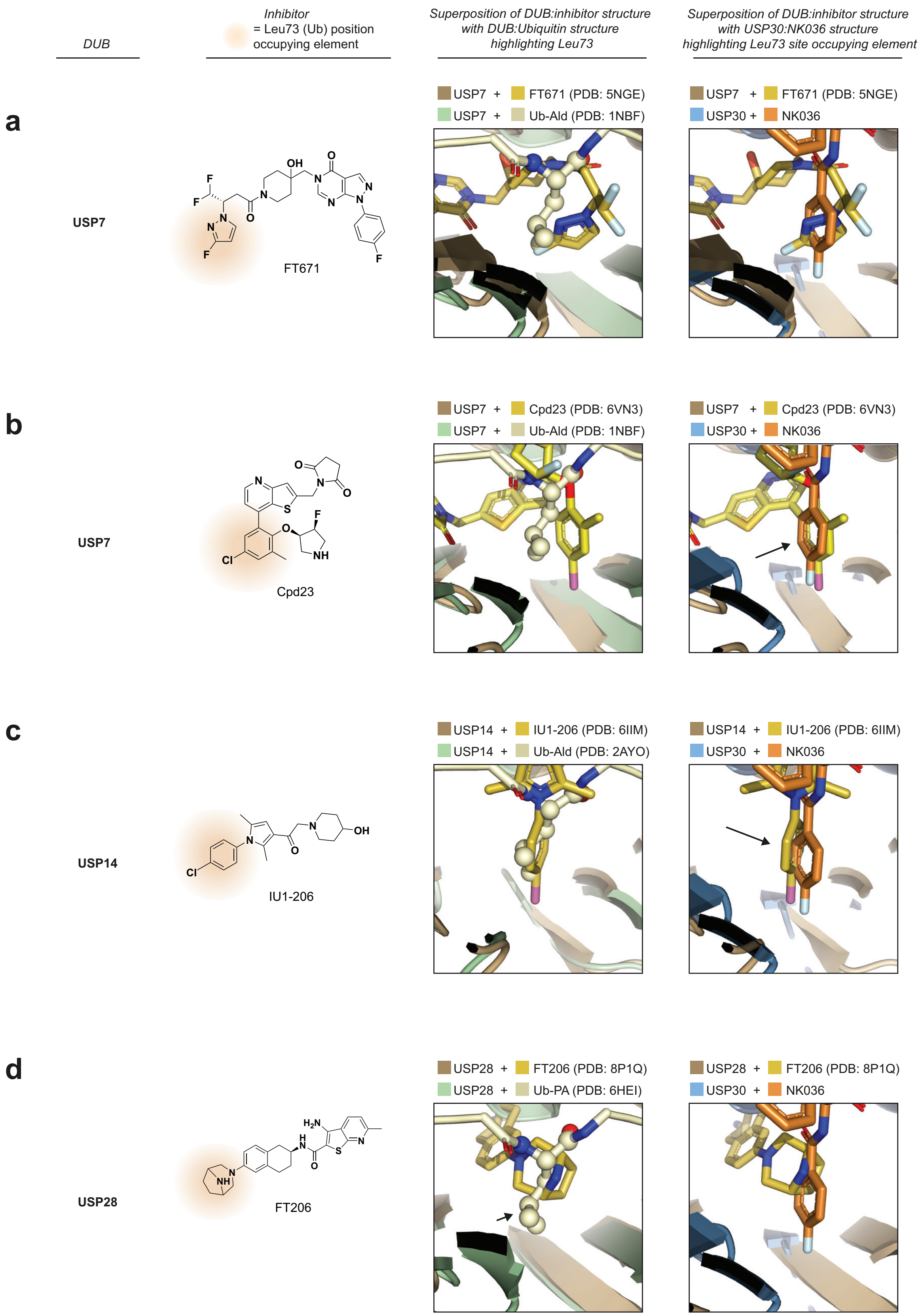
DUB
Inhibitor
= Leu73 (Ub) position occupying element
Superposition of DUB:inhibitor structure with DUB:Ubiquitin structure highlighting Leu73
Superposition of DUB:inhibitor structure with USP30:NK036 structure highlighting Leu73 site occupying element
a
USP7
FT671
USP7 + FT671 (PDB: 5NGE)
USP7 + Ub-Ald (PDB: 1NBF)
USP7 + FT671 (PDB: 5NGE)
USP30 + NK036
b
USP7
Cpd23
USP7 + Cpd23 (PDB: 6VN3)
USP7 + Ub-Ald (PDB: 1NBF)
USP7 + Cpd23 (PDB: 6VN3)
USP30 + NK036
c
USP14
IU1-206
USP14 + IU1-206 (PDB: 6IIM)
USP14 + Ub-Ald (PDB: 2AYO)
USP14 + IU1-206 (PDB: 6IIM)
USP30 + NK036
d
USP28
FT206
USP28 + FT206 (PDB: 8P1Q)
USP28 + Ub-PA (PDB: 6HEI)
USP28 + FT206 (PDB: 8P1Q)
USP30 + NK036

#### Supplementary Fig. 8 | A common DUB ligandability hotspot and a shared hotspot anchor.

**a-d,** Chemical and structural representations of inhibitors of USP7 (**a, b**), USP14 (**c**) and USP28 (**d**). Shown are the overlays of the inhibitor-bound structures with ubiquitin-bound structures of the respective DUBs (first box) and the inhibitor-bound structures of each DUB overlaid on the NK036 bound structure of USP30 (second box). Highlighted is the close-up view of the compound binding site at the ubiquitin Leu73 pocket. Chemically related hotspot anchors are indicated with black arrows. Structural superpositions were aligned taking only the protein residues into account. The chemical moieties of the inhibitors that occupy the Leu73 pocket are highlighted with orange background.

#### Supplementary Table 1 | Crystal structures of human USP family DUB catalytic domains.

| USP | $Zn^{2+}$ coordination at tip of fingers subdomain | PDB accession codes of | | | Comments |
|---|---|---|---|---|---|
| | | apo structures | Ub-bound structures | Inhibitor-bound structures | |
| USP1 | yes | 7AYO* | 7 AY2* | 7ZH4# (by Cryo-EM) | *in complex with UAF1 #with Ub-conjugated to FANCD2 |
| USP2 | yes | | 2HD5, 2IBI, 3NHE, 5XU8*, 5XVE, (+ with Ub variants) | | *in complex with Ub and 6-thioguanine |
| USP4 | yes | 2Y6E* | | | *insertions removed by limited proteolysis |
| USP5 | **no** | | 3IHP* | | *full-length |
| USP7 | **no** | 1NB8, 2F1Z, 4M5W, 4M5X, 5FWI, 5J7T | 1NBF, 5JTJ, 5JTV | 5N9R, 5N9T, 5NGE, 5NGF, 5UQV, 5UQX, 5VS6, 5VSB, 5VSK, 5WHC, 6F5H | |
| USP8 | yes | 2GFO | 3N3K* | | *with ubiquitin variant |
| USP9X | yes | 5WCH* | | | *with surface entropy reducing mutations |
| USP11 | yes | | 8OYP* | | *with Ub-GGG, insertion replaced by RDFrzS tag |
| USP12 | yes | 5K1A*, 5K1B*, 5K1C*#, 5K16 | 5L8W* | | *in complex with UAF1 #in complex with WDR20 |
| USP14 | **no** | 2AYN | 2AYO, 5GJQ* | 6IIN, 6IIM, 6IIL, 6IIK | *Cryo-EM structure of 26S proteasome |
| USP15 | yes | 6GHA | 6ML1*, 6CPM*, 6CRN* | 6GH9 | *with ubiquitin variant |
| USP21 | yes | | 2Y5B, 3I3T, 3MTN* | | *with ubiquitin variant |
| USP25 | **no** | 5O71, 6H4J, 6HEL | | | |
| USP28 | **no** | 6H4I, 6HEH*, 6HEJ, 8P19* | 6H4H, 6HEI*, 6HEK | 8HJE*, 7TUO*, 8P14*, 8P1P*, 8P1Q* | *insertions removed |
| USP30 | yes | | 5OHK#, 5OHN#, 9F6G* | 9F19*, 8D0A+, 8D1T+ | #insertions removed and with mutations *chimeric construct + with Fab |
| USP34 | yes | 7W3R | 7W3U | | |
| USP35 | yes | | 5TXK* | | *insertions removed |
| USP36 | yes | | 8BS9, (8BS3*) | | *with Fubi-PA |
| USP46 | yes | 6JLQ*# | 5CVO*, 5CVN*, 5CVM, 5L8H | | *in complex with UAF1 #in complex with WDR20 |
| CYLD | **no** | 2VHF | | | |

**Supplementary Table 2 | Biochemical and biophysical characterization of USP30 constructs.**

| **Construct** | **Catalytic efficiency** (x10$^3$ $M^{-1}s^{-1}$) | **Apo USP30 stability** (avg. $T_m$ in °C) | **Compound inhibitory potency** ($IC_{50}$ in nM) | | **Change in stability upon Inhibitor binding** (avg. $\Delta T_m$ in °C) | | **USP30 stability in presence of compound** (avg. $T_m$ in °C) | |
|---|---|---|---|---|---|---|---|---|
| | | | Cmpd39 | NK036 | Cmpd39 | NK036 | Cmpd39 | NK036 |
| c1 | 213.9 ± 2.4 | 51.1 | 0.8 ± 0.04 | 7.4 ± 0.3 | 7.8 | 7.2 | 58.9 | 58.3 |
| ch1 | 655.9 ± 17.6 | 46.5 | 0.7 ± 0.04 | 4.8 ± 1.7 | 9.6 | 8.7 | 56.1 | 55.2 |
| ch2 | 221.8 ± 3.6 | 50.7 | 0.5 ± 0.06 | 5.5 ± 0.6 | 8.4 | 7.8 | 59.1 | 58.5 |
| ch3 | 195.5 ± 2.4 | 54.9 | 0.3 ± 0.03 | 4.1 ± 0.3 | 6.6 | 6.5 | 61.5 | 61.4 |
| ch4 | 5.8 ± 0.0 | 51.5 | - | - | 8.8 | 7.7 | 60.3 | 59.2 |
| c2 | 11.3 ± 0.3 | 50.7 | - | - | 7.8 | 7.5 | 58.5 | 58.2 |
| ch3 F453Y | 62.7 ± 0.7 | 56.4 | 531 ± 99 | 632 ± 119 | 2.1 | 1.8 | 58.5 | 58.2 |
| ch3 L328F | 14.2 ± 0.2 | 57.0 | - | - | 1.5 | 1.2 | 58.5 | 58.2 |

#### Supplementary Table 3 | Statistics of individual datasets used for multi-crystal averaging.

| | USP30 + NK036 (crystal 1) | USP30 + NK036 (crystal 2) | USP30 + NK036 (crystal 3) | USP30 + NK036 (blended from crystals 1+2+3) |
|---|---|---|---|---|
| **Data collection** | | | | |
| Beamline | ESRF ID30A-3 | ESRF ID30A-3 | ESRF ID30A-3 | ESRF ID30A-3 |
| Wavelength | 0.9677 Å | 0.9677 Å | 0.9677 Å | 0.9677 Å |
| Space group | $P\,2_1\,2\,2_1$ | $P\,2_1\,2\,2_1$ | $P\,2_1\,2\,2_1$ | $P\,2_1\,2\,2_1$ |
| Cell dimensions | | | | |
| *a*, *b*, *c* (Å) | 55.90, 73.64, 201.66 | 55.96, 73.15, 200.37 | 55.92, 74.14, 201.67 | 55.83, 73.84, 201.05 |
| $\alpha, \beta, \gamma$ (°) | 90, 90, 90 | 90, 90, 90 | 90, 90, 90 | 90, 90, 90 |
| Anisotropy correction | yes | yes | yes | yes |
| Observed reflections | 59,699 | 49,129 | 62,471 | 214,419 |
| Unique reflections | 12,735 | 11,139 | 12,592 | 15,438 |
| Resolution (Å) | 48.89 – 2.95 (3.23 – 2.95) | 68.71 – 3.03 (3.37 – 3.03) | 53.89 – 3.01 (3.31 – 3.01) | 59.52 – 2.75 (3.23 – 2.75) |
| Ellipsoidal resolution limits (Å) [direction] | 2.95 [a*] 3.81 [b*] 2.95 [c*] | 2.97 [a*] 3.67 [b*] 3.28 [c*] | 2.95 [a*] 3.84 [b*] 3.01 [c*] | 2.89 [a*] 3.53 [b*] 2.75 [c*] |
| $R_{merge}$ | 0.131 (1.274) | 0.120 (0.729) | 0.134 (1.079) | 0.166 (1.536) |
| $R_{meas}$ | 0.147 (1.397) | 0.137 (0.888) | 0.150 (1.193) | 0.173 (1.595) |
| $I/\sigma(I)$ | 7.2 (1.4) | 6.6 (1.7) | 7.3 (1.4) | 9.6 (1.9) |
| $CC_{1/2}$ | 0.955 (0.611) | 0.997 (0.526) | 0.959 (0.631) | 0.998 (0.730) |
| Spherical completeness (%) | 69.7 (20.2) | 66.8 (14.9) | 72.8 (25.6) | 68.7 (26.2) |
| Ellipsoidal completeness (%) | 89.3 (56.6) | 85.2 (48.8) | 90.7 (62.1) | 91.9 (69.0) |
| Redundancy | 4.7 (5.4) | 4.4 (3.2) | 5.0 (5.6) | 13.9 (13.9) |
| Wilson *B* (Å$^2$) [direction] | 85 [a*] 166 [b*] 77 [c*] | 84 [a*] 177 [b*] 56 [c*] | 77 [a*] 158 [b*] 70 [c*] | 82 [a*] 161 [b*] 71 [c*] |

#### Protein sequences

Construct c1 – USP30Δ
*boundaries:* **64-178; GSGS; 217-357; SNA; 432-502**
*mutations:* **F348D, M350S, I353E**
KGLVPGLVNLGNTCFMNSLLQGLSACPAFIRWLEEFTSQYSRDQKEPPSHQYLSLTLLHLLKALSCQEVT
DDEVLDASCLLDVLRMYRWQISSFEEQDAHELFHVITSSLEDERDGSGSHWKSQHPFHGRLTSNMVCK
HCEHQSPVRFDTFDSLSLSIPAATWGHPLTLDHCLHHFISSESVRDVVCDNCTKIEAKGTLNGEKVEHQR
TTFVKQLKLGKLPQCLCIHLQRLSWSSHGTPLKRHEHVQFNE**D**L**S**MD**E**YKYHSNASTYLFRLMAVVVHH
GDMHSGHFVTYRRSPPSARNPLSTSNQWLWVSDDTVRKASLQEVLSSSAYLLFYERV

Construct ch1 – USP30 (USP7)
*boundaries:* **64-178; GSGS; 217-225 (326-348) 249-275 (370-398) 318-357; SNA; 432-502**
*mutations:* **F348D, M350S, I353E**
KGLVPGLVNLGNTCFMNSLLQGLSACPAFIRWLEEFTSQYSRDQKEPPSHQYLSLTLLHLLKALSCQEVT
DDEVLDASCLLDVLRMYRWQISSFEEQDAHELFHVITSSLEDERDGSGSHWKSQHPFHGKMVSYIQCKE
VDYRSDRREDYYDSLSLSIPAATWGHPLTLDHCLHHFISVEQLDGDNKYDAGEHGLQEAEKGVKFLTLPQ
CLCIHLQRLSWSSHGTPLKRHEHVQFNE**D**L**S**MD**E**YKYHSNASTYLFRLMAVVVHHGDMHSGHFVTYRR
SPPSARNPLSTSNQWLWVSDDTVRKASLQEVLSSSAYLLFYERV

Construct ch2 – USP30 (USP14)
*boundaries:* **64-178; GSGS; 217-224 (248-272) 249-275 (295-320) 318-357; SNA; 432-502**
*mutations:* **F348D, M350S, I353E**
KGLVPGLVNLGNTCFMNSLLQGLSACPAFIRWLEEFTSQYSRDQKEPPSHQYLSLTLLHLLKALSCQEVT
DDEVLDASCLLDVLRMYRWQISSFEEQDAHELFHVITSSLEDERDGSGSHWKSQHPFGVEFETTMKCTE
SEEEEVTKGKENQDSLSLSIPAATWGHPLTLDHCLHHFISQEEITKQSPTLQRNALYIKSSKISRLPQCLCI
HLQRLSWSSHGTPLKRHEHVQFNE**D**L**S**MD**E**YKYHSNASTYLFRLMAVVVHHGDMHSGHFVTYRRSPPS
ARNPLSTSNQWLWVSDDTVRKASLQEVLSSSAYLLFYERV

Construct ch3 – USP30 (USP14) [USP35]
*boundaries:* **64-178; GSGS; 217-224 (248-272) 249-275 (295-320) 318-348 [833-845] 437-502**
*mutations:* **F348D**
KGLVPGLVNLGNTCFMNSLLQGLSACPAFIRWLEEFTSQYSRDQKEPPSHQYLSLTLLHLLKALSCQEVT
DDEVLDASCLLDVLRMYRWQISSFEEQDAHELFHVITSSLEDERDGSGSHWKSQHPFGVEFETTMKCTE
SEEEEVTKGKENQDSLSLSIPAATWGHPLTLDHCLHHFISQEEITKQSPTLQRNALYIKSSKISRLPQCLCI
HLQRLSWSSHGTPLKRHEHVQFNE**D**LRLPLAGGRGQAYRLMAVVVHHGDMHSGHFVTYRRSPPSARN
PLSTSNQWLWVSDDTVRKASLQEVLSSSAYLLFYERV

Construct ch4 – USP30 (CYLD)
*boundaries:* **64-178; GSGS; 217-225 (700-713) 250-275 (742-746) 317-357; SNA; 432-502**
*mutations:* **F348D, M350S, I353E**
KGLVPGLVNLGNTCFMNSLLQGLSACPAFIRWLEEFTSQYSRDQKEPPSHQYLSLTLLHLLKALSCQEVT
DDEVLDASCLLDVLRMYRWQISSFEEQDAHELFHVITSSLEDERDGSGSHWKSQHPFHLKIRSAGQKVQ
DCYSLSLSIPAATWGHPLTLDHCLHHFISNLKFALPQCLCIHLQRLSWSSHGTPLKRHEHVQFNE**D**L**S**MD
**E**YKYHSNASTYLFRLMAVVVHHGDMHSGHFVTYRRSPPSARNPLSTSNQWLWVSDDTVRKASLQEVLS
SSAYLLFYERV

Construct c2 – USP30 (GS)
*boundaries:* **64-178; GSGS; 217-225; GSGS; 250-276; GSGSGS; 317-357; SNA; 432-502**
*mutations:* **F348D, M350S, I353E**
KGLVPGLVNLGNTCFMNSLLQGLSACPAFIRWLEEFTSQYSRDQKEPPSHQYLSLTLLHLLKALSCQEVT
DDEVLDASCLLDVLRMYRWQISSFEEQDAHELFHVITSSLEDERDGSGSHWKSQHPFHGSGSSLSLSIPA
ATWGHPLTLDHCLHHFISGSGSGSLPQCLCIHLQRLSWSSHGTPLKRHEHVQFNE**D**L**S**MD**E**YKYHSNAS
TYLFRLMAVVVHHGDMHSGHFVTYRRSPPSARNPLSTSNQWLWVSDDTVRKASLQEVLSSSAYLLFYE
RV

#### Supplementary Methods

General notes on synthetic methods

The chemicals and solvents used for this work were purchased from companies such as Activate Scientific, BLDpharm, Fisher Scientific, Sigma-Aldrich, TCI and VWR and used without further purification. Solvents used for the synthesis were named as follows. EA: ethyl acetate, DCM: dichloromethane, MeOH: methanol, DMF: dimethylformamide, DMSO: dimethyl sulfoxide, THF: tetrahydrofuran, ACN: acetonitrile, $H_2O$: water, EtOH: ethanol. Silica gel aluminum plates (silica gel 60 F254, Merck) were used for thin-layer chromatography and the detection was carried out using UV light at 254 and 360 nm. A Pure C-850 FlashPrep system (Büchi) was used for automated column chromatographic purification of the final product using a VP125/21 Nucleodur C18 Gravity column (21 x 125 mm; 5 µm, Macherey Nagel) by using a 0-100% gradient of $H_2O$ + 0.1% TFA to ACN + 0.1% TFA over 70 min.

A 1200 Infinity series HPLC system (Agilent Technologies) with a ZORBAX Eclipse XDB column (C18 80 Å; 4.6 x 150 mm; 5 µm, Agilent Technologies) was used for low resolution LC-MS analysis. For high resolution mass spectrometry (HRMS) an LTQ Orbitrap Fourier transform mass spectrometer (Thermo Fisher Scientific) coupled to an HPLC instrument from the same company (Hypersil Gold: 1 x 50 mm; 1.9 µm) was used. $H_2O$ + 0.1% formic acid and ACN + 0.1% formic acid were used as eluents. The ionization mode was ESI (Electrospray ionization) with a source voltage of 3.8 kV.

The following devices from Bruker were used to record NMR spectra: AV 500 Avance III HD (500 MHz for $^1$H and 125 MHz for $^{13}$C-NMR), AV 600 Avance III HD (600 MHz for $^1$H and 151 MHz for $^{13}$C NMR) and AV 700 Avance III HD (700 MHz for $^1$H and 176 MHz for $^{13}$C NMR). The chemical shifts of all spectra are specified in ppm and the coupling constants *J* are given in Hertz (Hz). Peaks of deuterated solvents were used as internal standards for $^1$H / $^{13}$C data (DMSO-$d_6$: δ = 2.50 ppm / 39.52 ppm). The multiplicities of the signals are abbreviated as follows: s (singlet), d (doublet), dd (doublet of doublets), ddd (doublet of doublets of doublets), t (triplet), q (quartet) and m (multiplet).

*Synthesis of (S)-4-fluoro-N-(1-((4-(N-(1-hydroxy-2-methylpropan-2-yl)sulfamoyl)phenyl)-amino)-1-oxo-3-phenylpropan-2-yl)benzamide (NK036)*

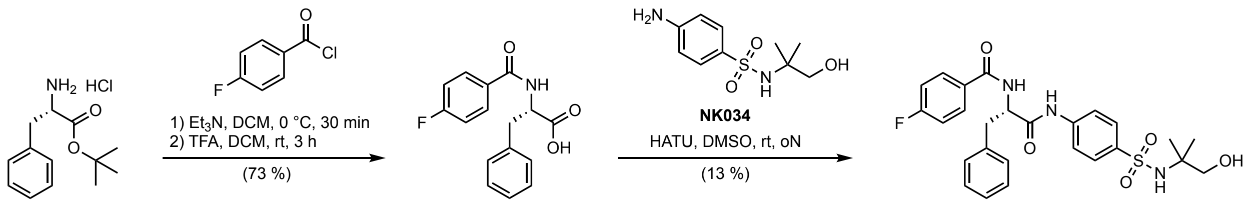

**NK020** **NK036**

The synthesis was based on previously described methods[33] for the synthesis of compound 39, with adjustments for the synthesis of (4-fluorobenzoyl)-L-phenylalanine (NK020) based on the synthesis of a related compound.[69] A different synthesis route of the title compound NK036 (I-137) was published elsewhere.[35]

Synthesis of (4-fluorobenzoyl)-L-phenylalanine (NK020)

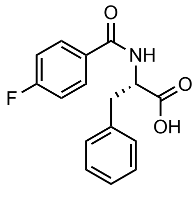

Step 1:
*tert*-Butyl L-phenylalaninate hydrochloride (2.8 g, 11 mmol, 1.0 eq) and triethylamine (2.8 g, 27.5 mmol, 2.5 eq) were dissolved in DCM (30 mL) and cooled to 0°C. 4-Fluorobenzoyl chloride (2.1 g, 13.2 mmol, 1.2 eq) was added dropwise while stirring. The reaction was stirred for 30 min at 0°C until completion. The reaction was quenched by the addition of water (50 mL). The aqueous layer was extracted with DCM (2 x 50 mL), and the combined organic phases were washed with brine (2 x 50 mL) and dried over MgSO4. The solvent was evaporated under reduced pressure to obtain the product *tert*-butyl (4-fluorobenzoyl)-L-phenylalaninate (NK019) as a yellow oil. The crude product was used for the next step without further purification.

Step 2:
*tert*-Butyl (4-fluorobenzoyl)-L-phenylalaninate (3.7 g, 11 mmol) was dissolved in HCl/dioxan (4 M, 20 mL) and stirred at rt for 3 h. After completion, the solvents were evaporated under reduced pressure to obtain the product (4-fluorobenzoyl)-L-phenylalanine (NK020) as an off-white powder (2.9 g, 10.1 mmol, 73 % over two steps).

**$^1$H NMR** (600 MHz, DMSO) δ (ppm): 12.81 (s, 1H), 8.77 (d, *J* = 8.2 Hz, 1H), 7.89 (dd, *J* = 8.8, 5.6 Hz, 2H), 7.34 – 7.31 (m, 2H), 7.31 – 7.24 (m, 4H), 7.20 – 7.15 (m, 1H), 4.65 (ddd, *J* = 10.7, 8.1, 4.4 Hz, 1H), 3.22 (dd, *J* = 13.8, 4.4 Hz, 1H), 3.08 (dd, *J* = 13.9, 10.7 Hz, 1H).

**$^{13}$C NMR** (151 MHz, DMSO) δ (ppm): 173.25, 165.42, 164.03 (d, *J* = 248.7 Hz), 138.21, 130.42 (d, *J* = 2.8 Hz), 130.08 (d, *J* = 8.9 Hz), 129.10, 128.26, 126.43, 115.25 (d, *J* = 21.7 Hz), 54.35, 36.34.

**HRMS** m/z for $C_{16}H_{15}FNO_3^+$ ($[M+H]^+$) calculated: 288.1031, found: 288.1029.

Synthesis of (*S*)-4-fluoro-*N*-(1-((4-(*N*-(1-hydroxy-2-methylpropan-2-yl)sulfamoyl)phenyl)-amino)-1-oxo-3-phenylpropan-2-yl)benzamide (NK036)

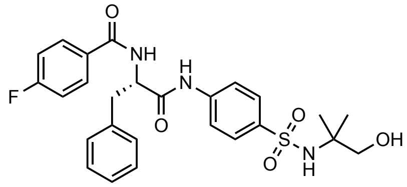

4-Fluorbenzoyl-L-phenylalanine (NK020) (70.6 mg, 0.25 mmol, 1.2 eq) was dissolved in dry DMSO (2 mL). To that solution was added HATU (155.6 mg, 0.41 mmol, 2 eq) and the resulting mixture was stirred for 30 minutes at rt. 4-Amino-*N*-(1-hydroxy-2-methylpropan-2-yl)benzenesulfonamide (NK034) (50 mg, 0.2 mmol, 1.0 eq) was added and the reaction stirred overnight. The resulting solution was then directly purified by reverse phase column chromatography. The product NK036 was obtained as a white powder (13.2 mg, 0.03 mmol, 13 %).

**$^1$H NMR** (700 MHz, DMSO-$d_6$) δ (ppm): 10.58 (s, 1H), 8.86 (d, *J* = 7.9 Hz, 1H), 7.92 – 7.89 (m, 2H), 7.77 (s, 4H), 7.41 (d, *J* = 7.1 Hz, 2H), 7.32 – 7.26 (m, 4H), 7.21 – 7.16 (m, 2H), 4.84 (ddd, *J* = 10.4, 7.9, 4.8 Hz, 1H), 3.17 (s, 2H), 3.16 – 3.08 (m, 2H), 1.00 (s, 6H).

**$^{13}$C NMR** (176 MHz, DMSO-$d_6$) δ (ppm): 171.08, 165.56, 163.97 (d, *J* = 248.8 Hz), 141.82, 138.59, 137.97, 130.22 (d, *J* = 2.9 Hz), 130.17 (d, *J* = 9.0 Hz), 129.17, 128.12, 127.40, 126.41, 118.87, 115.16 (d, *J* = 21.7 Hz), 68.93, 56.81, 56.06, 36.90, 24.02.

**HRMS** m/z for $C_{26}H_{29}FN_3O_5S^+$ ($[M+H]^+$) calculated: 514.1806, found: 514.1811.

*Synthesis of 4-amino-N-(1-hydroxy-2-methylpropan-2-yl)benzenesulfonamide (NK034)*

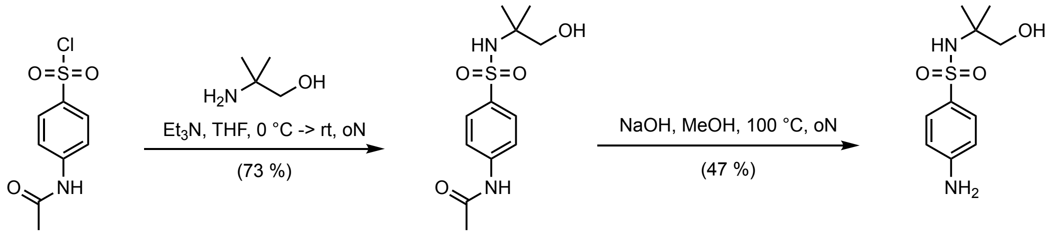

**NK027** **NK034**

Synthesis of *N*-(4-(*N*-(1-hydroxy-2-methylpropan-2-yl)sulfamoyl)phenyl)acetamide (NK027)

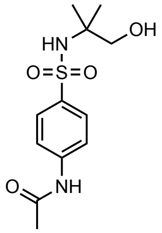

To a solution of 4-acetamidobenzenesulfonyl chloride (5 g, 21.4 mmol, 1.0 eq) in dry THF (30 mL) was added dropwise triethylamine (4.3 g, 42.8 mmol, 2 eq) and 2-amino-2-methyl-1-propanol (3.8 g, 42.8 mmol, 2 eq) at 0°C while stirring. The reaction was then warmed to rt and stirred over night until completion. The mixture was then poured into a mixture of ice and water (50 mL). The product was extracted with EA (3 x 50 mL) and the combined organic layers were washed with saturated $NH_4Cl$ solution (2 x 100 mL), water (1 x 100 mL) and brine (1 x 100 mL). The organic layer was dried over anhydrous $MgSO_4$ after which the solvents were removed under reduced pressure. The product *N*-(4-(*N*-(1-hydroxy-2-methylpropan-2-yl)sulfamoyl)phenyl)-acetamide (NK027) was obtained as a white solid (4.5 g, 15.7 mmol, 73 %).

**$^1$H NMR** (500 MHz, DMSO-$d_6$) δ (ppm): 10.28 (s, 1H), 7.81 – 7.65 (m, 4H), 7.17 (s, 1H), 4.73 (t, *J* = 5.9 Hz, 1H), 3.16 (d, *J* = 6.0 Hz, 2H), 2.08 (s, 3H), 0.98 (s, 6H).

**$^{13}$C NMR** (126 MHz, DMSO-$d_6$) δ (ppm): 168.96, 142.29, 138.13, 127.40, 118.43, 68.96, 56.81, 24.15, 24.03.

**LCMS** m/z for $C_{12}H_{19}N_2O_4S^+$ ($[M+H]^+$) calculated: 287.1, found: 287.2.

Synthesis of 4-amino-*N*-(1-hydroxy-2-methylpropan-2-yl)benzenesulfonamide (NK034)

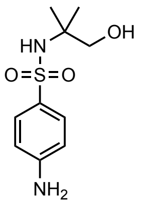

To a solution of *N*-(4-(*N*-(1-hydroxy-2-methylpropan-2-yl)sulfamoyl)phenyl) acetamide (NK027) (520 mg, 1.82 mmol) in methanol (4 mL) was added aq. NaOH (5 M, 3 mL). The reaction mixture was stirred for 100°C over night. After completion, the reaction was cooled to room temperature and diluted with water (10 mL). The organic solvent was removed under reduced pressure. The aqueous solution was then neutralized with aq. HCl (2 M) to pH 7-8, after which the product was extracted with EA (2 x 50 mL). The combined organic layers were dried over $MgSO_4$, and the solvent was removed under reduced pressure which yielded the product 4-amino-*N*-(1-hydroxy-2-methylpropan-2-yl)benzenesulfonamide (NK034) as a white solid (210 mg, 0.83 mmol) which was directly used for the next step without further purification.

**HRMS** m/z for $C_{10}H_{17}N_2O_3S^+$ ($[M+H]^+$) calculated: 245.0954, found: 245.0953.

*Synthesis of (S)-N-(1-((4-(N-(tert-Butyl)sulfamoyl)phenyl)amino)-1-oxo-3-phenylpropan-2-yl)-4-fluorobenzamide (compound 39)*

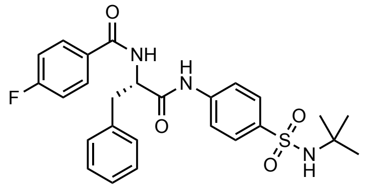

Compound 39 was synthesized using the same synthetic procedure as described above. The analytical data are in full agreement with previously published $^1$H NMR data.

**$^1$H NMR** (700 MHz, DMSO-$d_6$) δ (ppm): 10.58 (s, 1H), 8.86 (d, *J* = 7.9 Hz, 1H), 7.90 (dd, *J* = 8.7, 5.6 Hz, 2H), 7.77 (s, 4H), 7.42 – 7.40 (m, 3H), 7.29 (q, *J* = 8.1, 7.3 Hz, 4H), 7.18 (t, *J* = 7.4 Hz, 1H), 4.87 – 4.81 (m, 1H), 3.19 – 3.05 (m, 2H), 1.08 (s, 9H).

**$^{13}$C NMR** (176 MHz, DMSO-$d_6$) δ (ppm): 171.08, 165.55, 163.97 (d, *J* = 248.9 Hz), 141.81, 138.43, 137.96, 130.22 (d, *J* = 2.9 Hz), 130.17 (d, *J* = 9.0 Hz), 129.17, 128.12, 127.44, 126.40, 118.90, 115.16 (d, *J* = 21.8 Hz), 56.06, 53.11, 36.90, 29.73.

**HRMS** m/z for $C_{26}H_{29}FN_3O_4S^+$ ($[M+H]^+$) calculated: 498.1857, found 498.1850.

#### NMR spectra of compounds

NK020 $^{1}H$ NMR

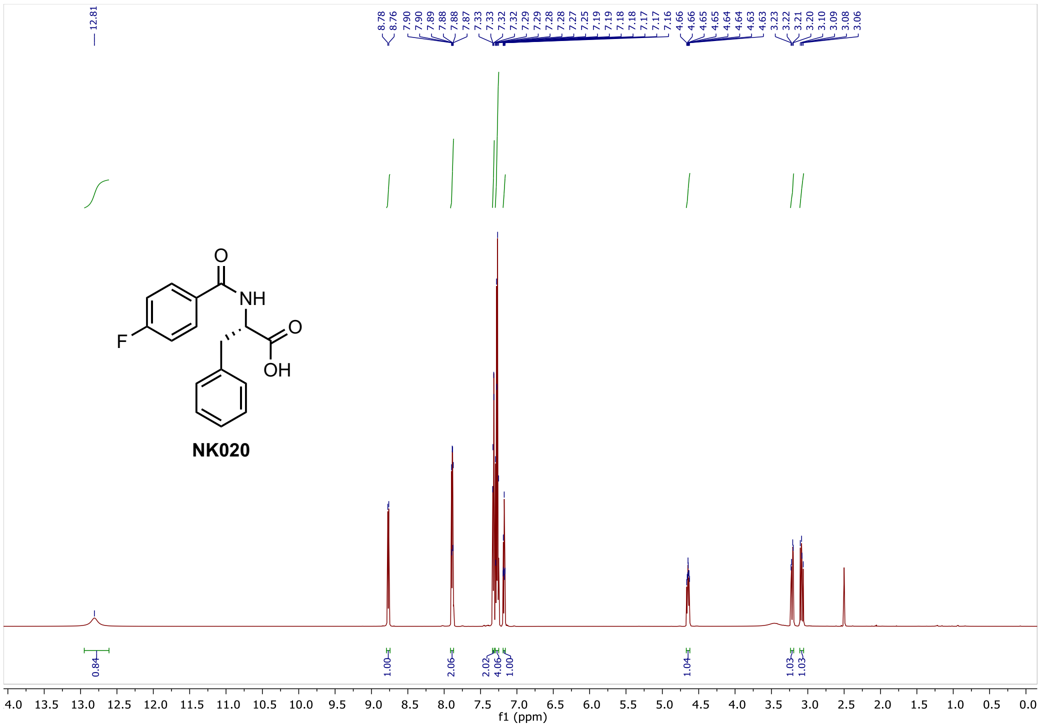

NK020 $^{13}C$ NMR

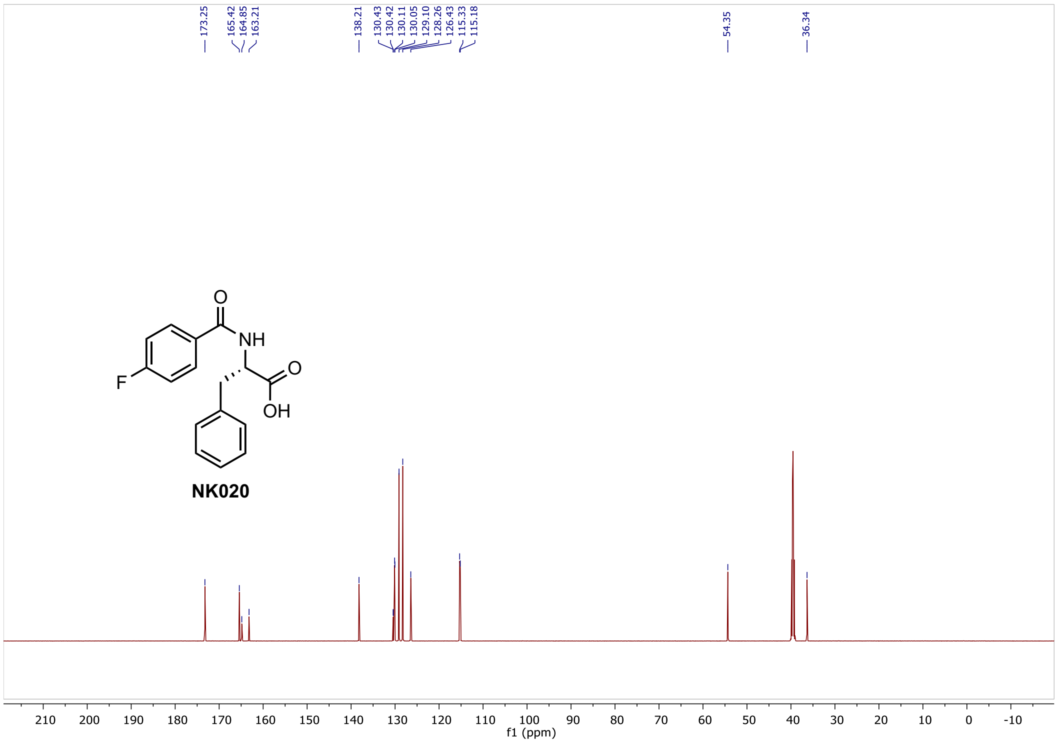

NK036 $^{1}$H NMR

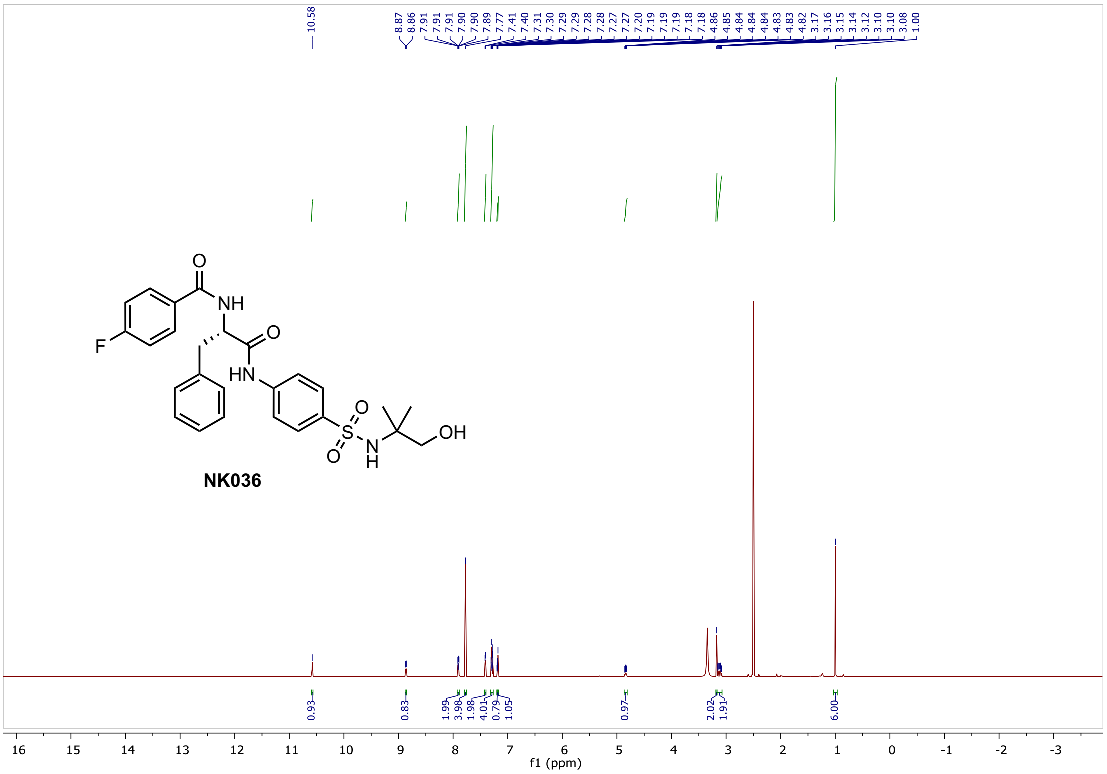

NK036 $^{13}$C NMR

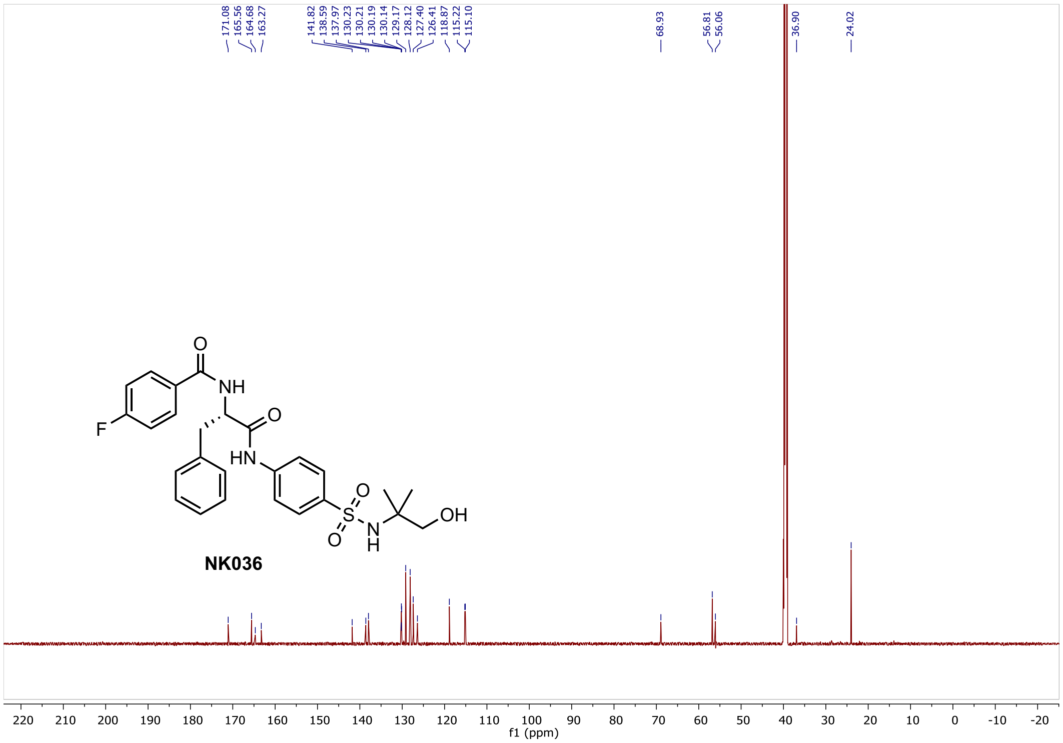

#### NK027 $^1$H NMR

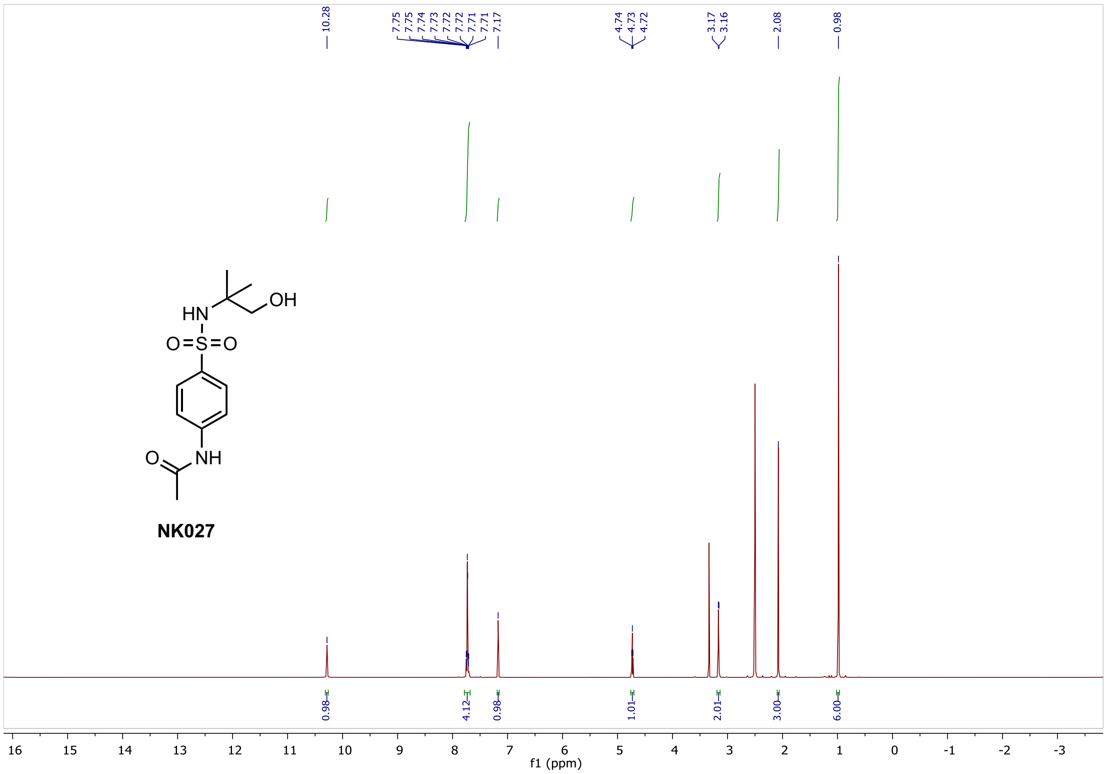

#### NK027 $^{13}$C NMR

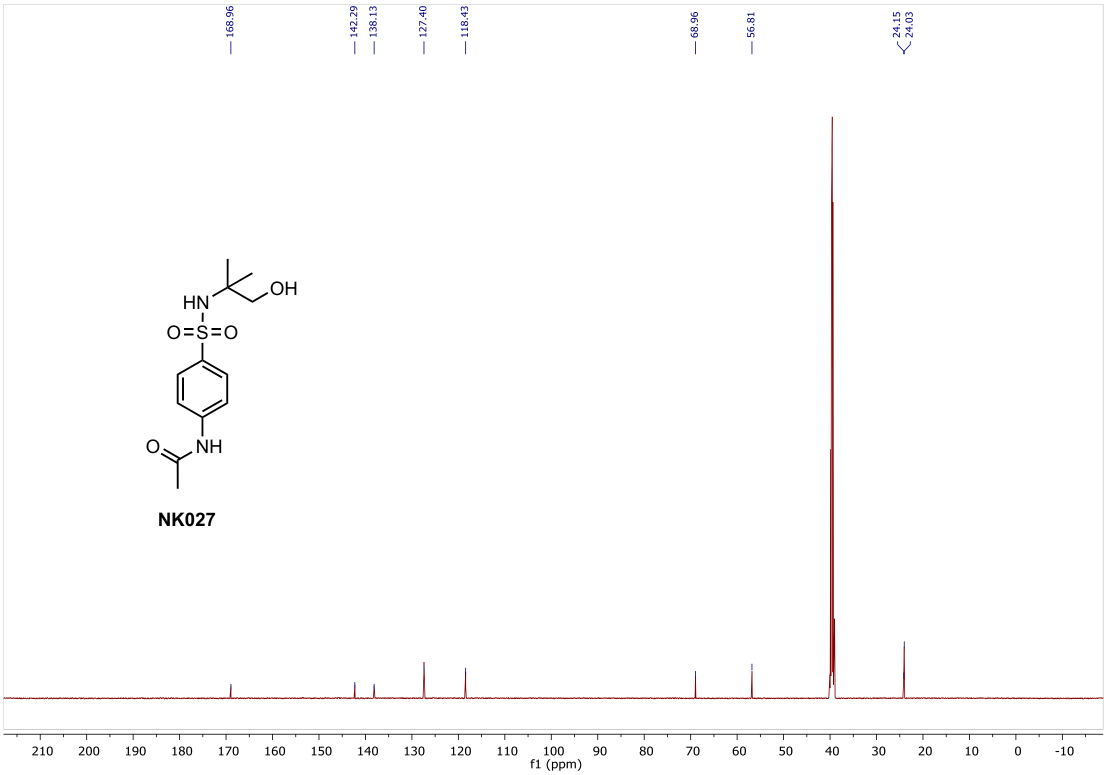

#### Compound 39 $^{1}$H NMR

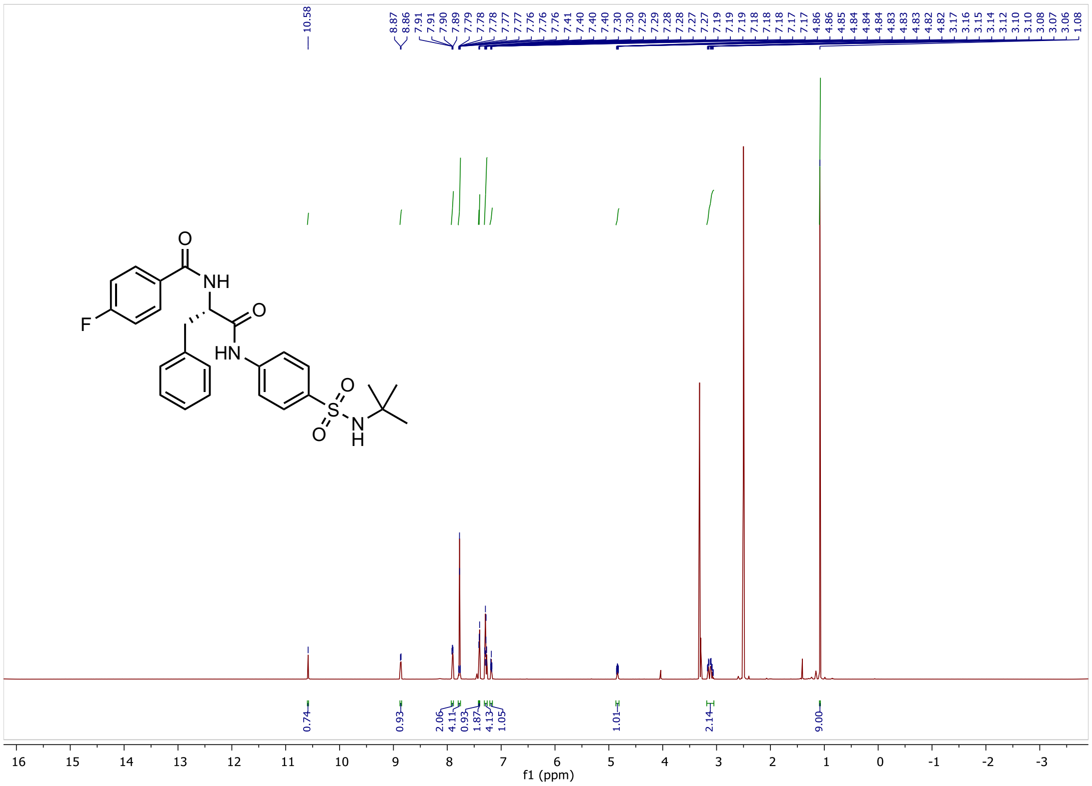

#### Compound 39 $^{13}$C NMR

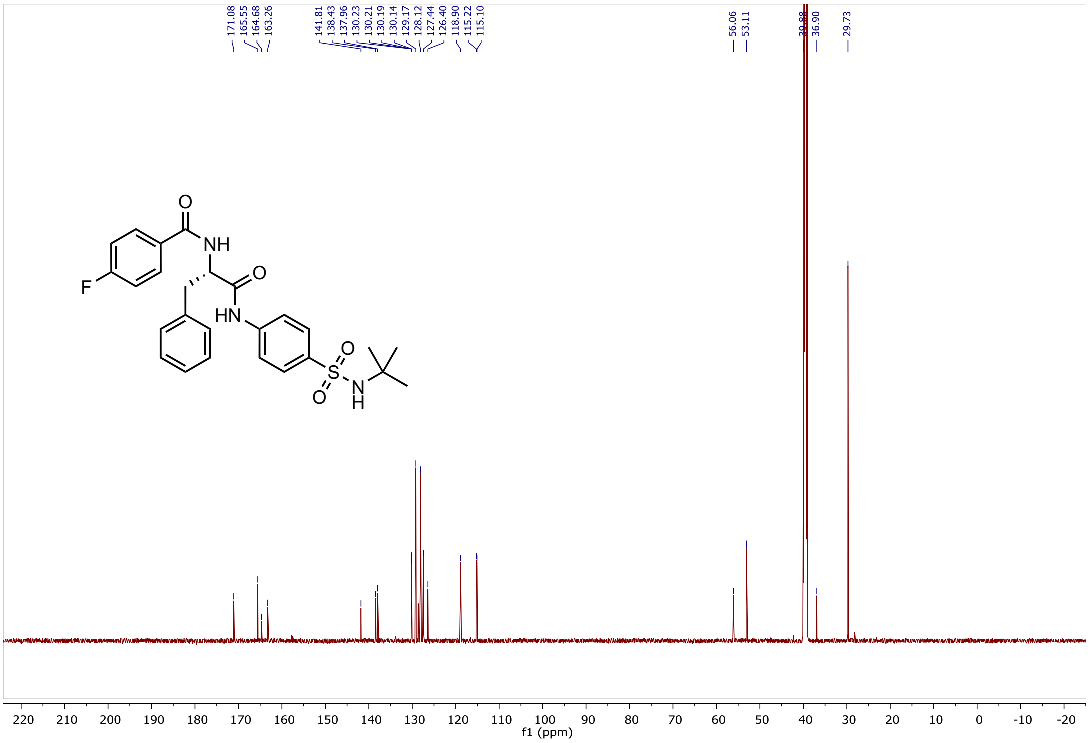

### Uncropped gels

Figure 1i

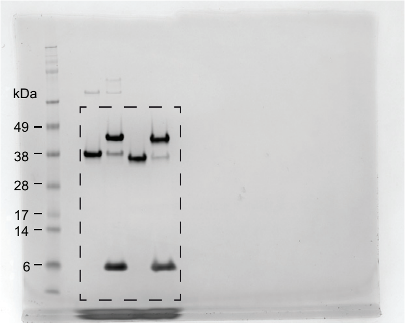

Supplementary Figure 3b

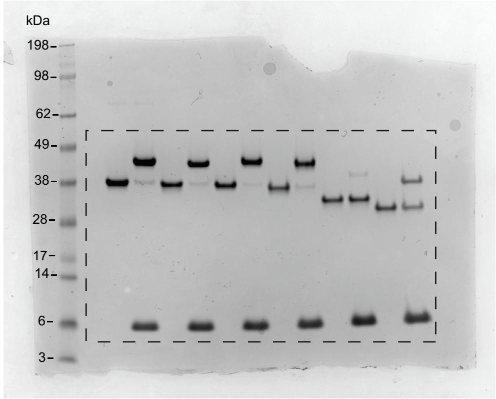

Figure 4i

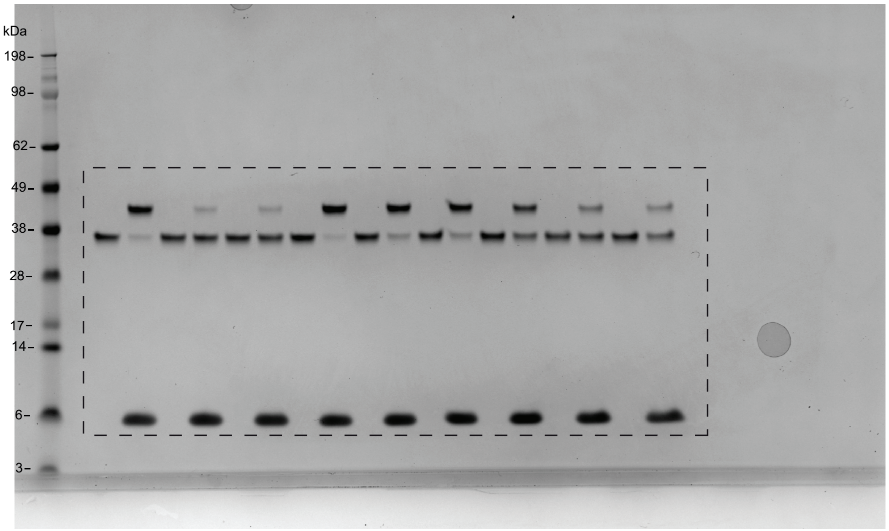

#### Uncropped blots

Figure 4j

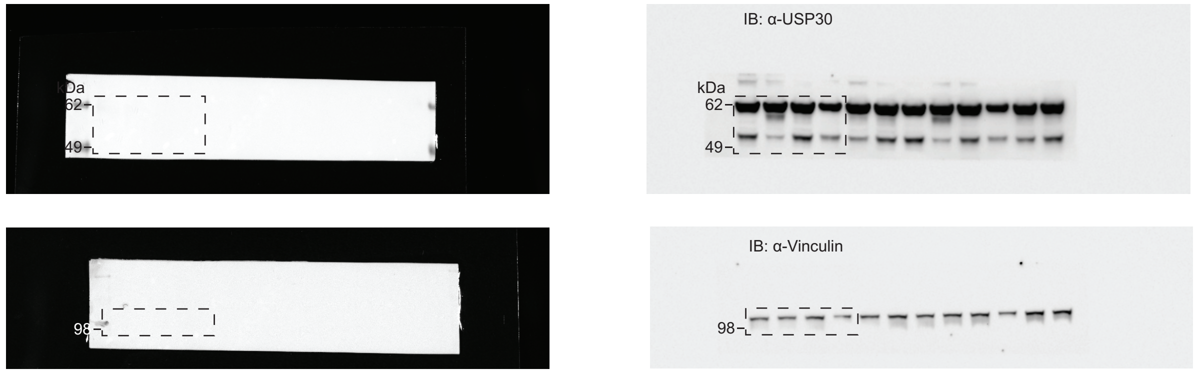

Figure 4k

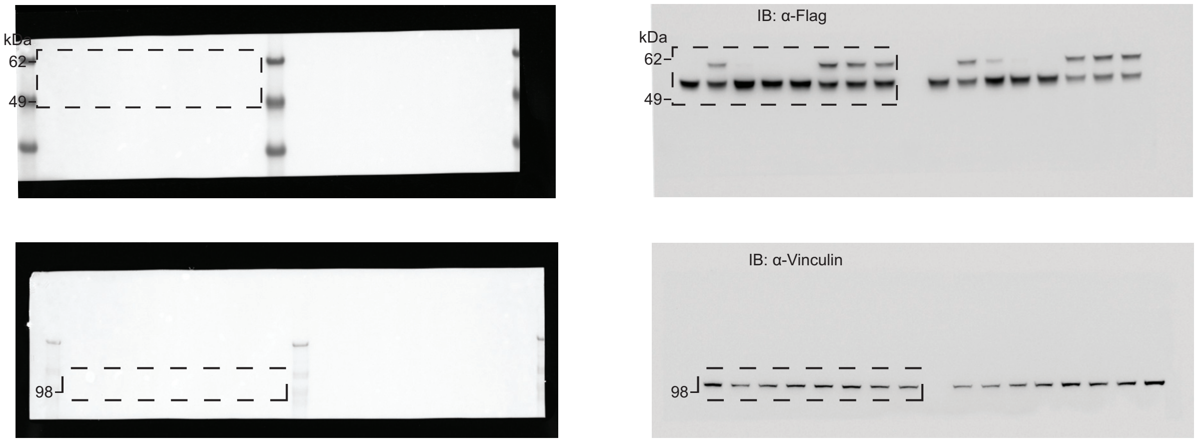
